# The highly conserved aphid effector pair Mp1-Mp58 associates to form an effector complex that targets host trafficking protein VPS52

**DOI:** 10.1101/2025.05.12.653458

**Authors:** Jade R. Bleau, Namami Gaur, S. Ronan Fisher, Thomas Waksman, Michael Porter, Jorunn IB Bos

## Abstract

Pathogen and pest effectors play a crucial role in manipulating plant biological processes, facilitating infection and infestation. While pathogens and pests secrete repertoires of effectors into their host plants, most effector function studies focus on characterising individual proteins. We previously identified a genetically linked and co-regulated gene pair in the aphid *Myzus persicae* encoding effectors Mp1 and Mp58. Here, we explored the functional link between these effectors. We used ectopic expression assays in *Nicotiana benthamiana* followed by co-immunoprecipitation assays and confocal microscopy to explore effector-effector and effector-target interactions and their subcellular localisation. We produced recombinant proteins to validate effector interactions and used computational modelling to predict effector complex 3D structures. We revealed that effectors Mp1 and Mp58 interact *in planta* and *in vitro* and likely form an oligomeric complex. Both effectors associate with the host target Vacuolar Protein Sorting associated Protein 52 (VPS52) to form an Mp1-Mp58-VPS52 complex which localises at vesicle-like structures. Our findings point to effector complex formation in plant-insect interactions and highlight a further layer of complexity in the molecular dialogue between plants and insects. Our work also shows the importance of considering the context in which effectors may function within a larger effector repertoire.

## Introduction

Phloem-feeding insects, such as plant hoppers, leaf hoppers, whiteflies, and aphids deliver molecules inside their host to promote susceptibility, known as effectors. Effector delivery takes place during phloem-feeding and probing and involves the secretion of saliva via highly specialised mouthparts called stylets. Saliva, containing effectors, is secreted into the apoplast, phloem and, depending on insect species, may also be delivered into cells along the stylet pathway (Naalden et al., 2021; Wang et al., 2023). Like plant pathogen effectors, which have extensively been studied over the past decades, phloem-feeding insect effectors target host proteins to modify their activity to promote host susceptibility, extending the effector paradigm to herbivorous insects. Advances in genomics and proteomics approaches have indeed unveiled effector repertoires of many herbivorous insect species, and characterisation of their virulence activities remains a major challenge. With limited genetic crop resistance available against insect pests, understanding how effectors function and interfere with plant host cell biology promises to underpin the design of novel strategies for crop protection.

The aphid *Myzus persicae* (green peach aphid) is a major agricultural pest in part due to its exceptionally broad host range and ability to vector many important plant viruses (Blackman and Eastop, 2000). The *M. persicae* effector repertoire consists of diverse proteins, as predicted by saliva proteomics and bioinformatics pipelines (Bos et al., 2010; Harmel et al., 2008; Thorpe et al., 2018, 2016; Vandermoten et al., 2014) and features an overrepresentation of disordered proteins (Waksman et al., 2024). Functional characterisation of *M. persicae* effector proteins led to the identification of several host targets (Gravino et al., 2024; Liu et al., 2024; Rodriguez et al., 2017; Wang et al., 2021). The *M. persicae* effector Mp1 is expressed in salivary glands, and secreted in saliva as shown by proteomics (Bos et al., 2010; Harmel et al., 2008; Thorpe et al., 2016). We previously showed that Mp1 can associate with Vacuolar Protein Sorting associated Protein 52 (VPS52) in a species-specific manner. Specifically, Mp1 was only able to interact with VPS52 proteins from host plant species *Solanum tuberosum* (StVPS52) and Arabidopsis (AtVPS52), but not with those from non/poor-host plants *Medicago truncatula* (MtVPS52) and *Hordeum vulgare* (HvVPS52). In line with these observations, Mp1 variants from aphid species unable to infest *S. tuberosum* and *Arabidopsis* did not interact with VPS52 from these plant species (Rodriguez et al., 2017). Over-expression of StVPS52 reduced host susceptibility to aphids, and infestation led to a reduction of detectable VPS52 protein levels, indicating that VPS52 is an important virulence target. Targeting of host cellular trafficking machinery by plant parasites is a common feature in plant-pathogen interactions (Yuen et al., 2023) and the association of aphid effector Mp1 with VPS52, a component of the Golgi Associated Retrograde Protein (GARP) complex, and degradation of VPS52 during infestation, indicate this feature can be extended to plant-herbivorous insect interactions.

The *Mp1* effector gene is co-located with another effector gene, *Mp58*, within the *M. persicae* genome, and the expression of this effector gene pair is tightly co-regulated (Thorpe et al., 2018). The effector genes are positioned in a head to tail orientation, around 5.5kb apart, and feature a highly similar 5’ end promoter region. Moreover, the co-location of this gene pair is conserved across at least five aphid genomes but with a lack of synteny in the corresponding genomic regions. Aphid gene expression analyses revealed tight co-regulation of Mp1(-like) and Mp58(-like) in different species as well as co-regulation with a set of putative effectors, pointing to a shared transcriptional control mechanism (Thorpe et al., 2018).

The effector Mp58, like Mp1, can be detected in saliva by proteomics, and is highly expressed in *M. persicae* salivary glands (Harmel et al., 2008; Thorpe et al., 2016). Ectopic expression of this effector in either Arabidopsis, *Nicotiana tabacum*, or *Nicotiana benthamiana* leads to reduced susceptibility to *M. persicae*, suggesting this effector may trigger plant defence responses (Elzinga and Jander, 2014; Escudero-Martinez et al, 2020). In contrast, the Mp58-like effector from *Macrosiphum euphorbiae*, called Me10, enhances tomato and *N. benthamiana* susceptibility to *M. euphorbiae* and *M. persicae* (Atamian et al., 2013), and interacts with tomato 14-3-3 isoform 7 (TFT7), which contributes to defence against aphids (Chaudhary et al., 2019).

The co-location and coregulation of the highly conserved *Mp1*-*Mp58* gene pair across aphid species point to a functional link between the encoded effector proteins. In this study we sought to explore this link using protein-protein interaction assays, confocal microscopy and computational modelling. We show that Mp1 and Mp58 physically interact to form an effector complex, and that both effectors can associate with VPS52, a previously identified Mp1 host target implicated in plant-aphid interactions. Computational modelling pointed to the N-terminal region of VPS52 encompassing the first 170 amino acids of the protein, and forming an α-helix, to be the site of interaction of Mp1. Prediction of the Mp58 interaction site pointed to a larger and less defined region of VPS52 being involved. We experimentally confirmed the interaction of Mp1 with the VPS52 N-terminus *in vitro* using the VPS52 1-170 domain and in planta using host/nonhost VPS52 chimeras. Moreover, in co-expression assays with individual effector proteins Mp1 re-localises to VPS52-associated vesicles, while Mp58 maintains cytoplasmic. However, when both effectors are co-expressed with VPS52, Mp58 can be detected together with Mp1 and VPS52 in vesicle-like structures pointing to the presence of an Mp58-Mp1-VPS52 protein complex. Our data highlight an unexplored layer of complexity in the plant-herbivorous insect molecular dialogue based on effector complex formation not only with host proteins but also with other effectors to potentially mediate and/or regulate virulence activities. Our study also highlights the importance of considering effector functions not in isolation, but in the context of secreted effector repertoires, with some effectors potentially mediating and/or regulation activities via interaction with other effectors.

## Results

### *Myzus persicae* effectors Mp1 and Mp58 physically interact to form an effector complex

We previously showed that the effector genes encoding Mp1 and Me10-like (Mp58 in *M. persicae*) are paired and genetically linked across the genomes of 5 aphid species, and feature shared transcriptional control (Thorpe et al., 2018). These observations led us to hypothesise that effectors Mp1 and Mp58 may function together during aphid infestation, potentially through protein-protein interaction. We first tested whether Mp1 and Mp58 can associate in planta by co-expressing myc-Mp1 with GFP-Mp58 in *Nicotiana benthamiana* followed by co-immunoprecipitation (co-IP). We were able to detect GFP-Mp58 in the myc-Mp1 immunoprecipitation, but not in the myc-GUS control, indicating that Mp1 and Mp58 associate (Fig 1A). To determine whether the effector pair interact directly or whether the interaction may be mediated by interacting plant proteins, we produced recombinant GST-Mp1 and His-Mp58 proteins in *Escherichia coli.* We individually expressed the proteins in *E. coli* and combined lysates before immobilized metal affinity chromatography (IMAC) of His-Mp58. Eluates were analysed by western blotting and showed co-purification of GST-Mp1, pointing to direct interaction between Mp1 and Mp58 (Fig. 1B).

**Figure 1:**
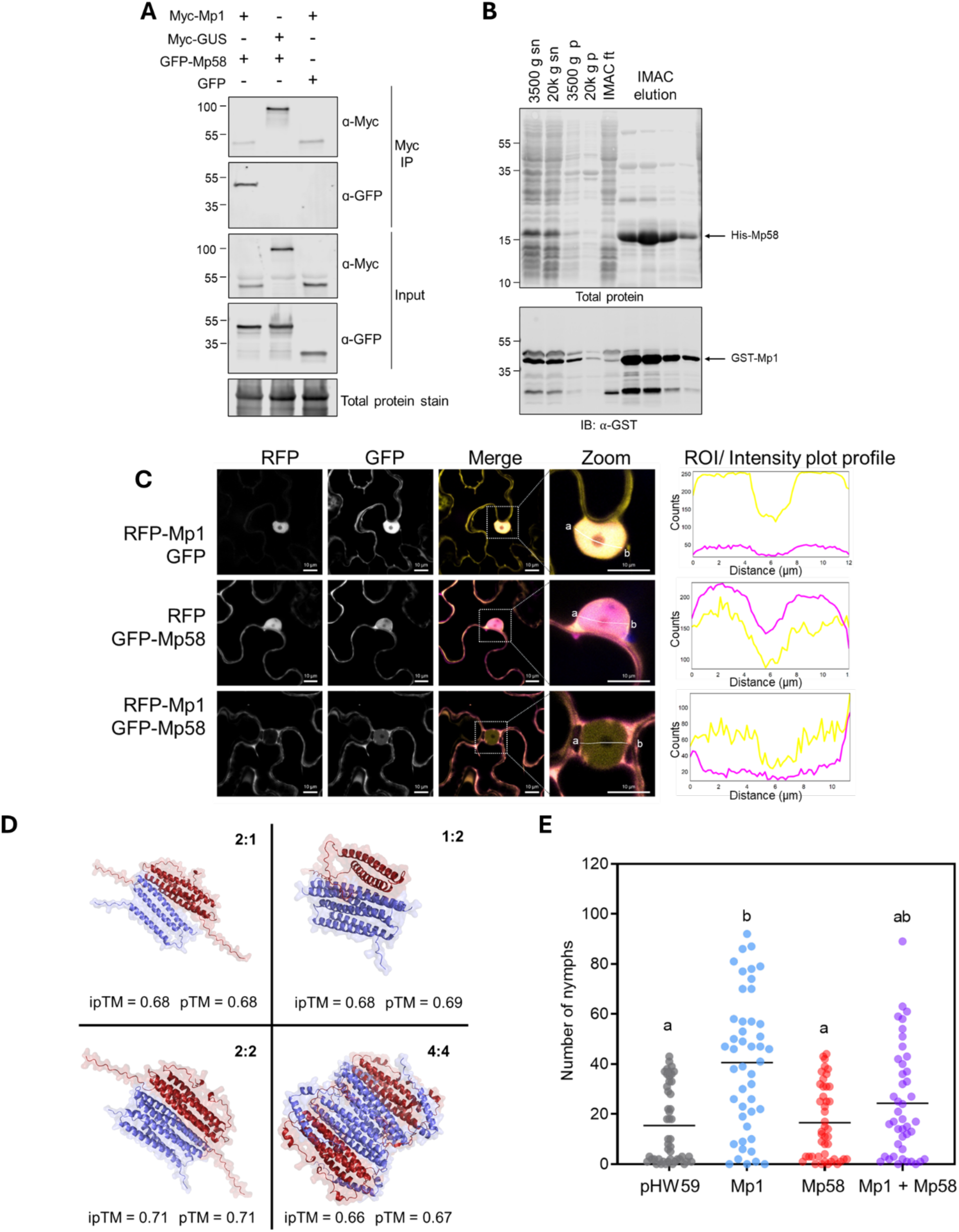
Mp1 and Mp58 interact *in vitro* and *in planta.* **(A)** Myc-Mp1 and GFP-Mp58 were transiently expressed via Agroinfiltration with each other or GFP and myc-GUS as controls. Protein were immunoprecipitated with myc-trap. **(B)** co-purification of His-Mp58 and GST-Mp1 from recombinant protein expressed in *E. coli*. Expression of His-Mp58 and GST-Mp1 was induced in *E.coli* individually and protein lysates were combined. Protein was purified via IMAC and blotted against GST. Sn - supernatant; p – pellet; ft – flowthrough; IMAC – immobilised metal affinity chromatography. Ladders represent kDa. **(C)** Co-localisation of RFP-Mp1 (in magenta) and GFP-Mp58 (in yellow) and GFP or RFP, respectively by confocal microscopy. Images were taken 3 dpi after agroinfiltration of *Nicotiana benthamiana*. Merged figured represent overlay images of the GFP and RFP channels. White boxes in merge panels indicate zoomed section. Scale bar = 10 µm, magnification 40X. Presented images are single plane images. **(D)** AlphaFold3 structural predictions for Mp1–Mp58 hetero-oligomeric states: Predictions were conducted using AF3 by manually by setting Mp1:Mp58 stoichiometric ratios within a range of all possible permutations from 1:1 to 4:4 and automatically using MultiFOLD2 (supplementary materials). Shown are the four oligomeric combinations of Mp1 (red) and Mp58 (blue) with the highest ipTM scores as assessed by AlphaFold3, where Mp1:Mp58 in a 2:1 ratio resulted in an ipTM of 0.68 and pTM of 0.68, 1:2 ratio ipTM: 0.68, pTM: 0.69, 2:2 ratio ipTM: 0.71, pTM: 0.71 and 4:4 ratio ipTM: 0.66, pTM 0.67. DeepUMQA-X analysis results in an Interface-lDDT of 0.626 for 2:1, 0.575 for 1:2, 0.652 for 2:2 and 0.656 for 4:4 **(E)** *N. benthamiana* leaves transiently expressing Mp1 and Mp58 under the phloem-specific promoter (AtSUC2) individually or together were challenged with *M. persicae* and fecundity was assessed over 14 days. Empty vector (pHW59) was used as a control. Empty vector pHW59 was included in individual effector samples (Mp1/Mp58 alone) to ensure the OD was consistent in all conditions. The graph shows all data points collected over three independent replicates (n = 9-12). The horizontal line represents the mean. A Kruskal-Wallis test and Dunn’s multiple comparisons test was conducted for statistical analysis of differences between groups. Different letters represent *p* < 0.01.

We previously performed subcellular localisation of GFP-Mp1 and GFP-Mp58, which showed a cytoplasmic/nuclear localisation for both proteins (Escudero-Martinez et al., 2020; Rodriguez et al., 2017). While we initially generated a mRFP-Mp58 construct for co-localisation experiments, we were unable to consistently detect RFP-Mp58 by western blotting and confocal microscopy, while, in line with previous observations, GFP-Mp58 was easily detectable. Therefore, we performed co-localisation experiments with the GFP-Mp58 and RFP-Mp1 combination. When individually expressed, RFP-Mp1 localises predominantly to the nucleus, with a weaker signal observed in the cytoplasm (Fig. 1C). The weak RFP-Mp1 signal observed in the cytoplasm compared to GFP-Mp1 in previous reports (Escudero-Martinez et al., 2020; Rodriguez et al., 2017) is possibly due to lower expression of RFP-Mp1 compared to GFP-Mp1, though we were able to detect RFP-Mp1 by western blotting and detect an interaction between RFP-Mp1 and GFP-Mp58 via co-IP (Fig. S1). GFP-Mp58, when expressed alone, localised to both the nucleus and cytoplasm (Fig. 1C) as previously described (Escudero-Martinez et al., 2020). When RFP-Mp1 and GFP-Mp58 were co-expressed, GFP-Mp58 remained localised to the nucleus and the cytoplasm, whereas RFP-Mp1 was no longer present in the nucleus and only detected in the cytoplasm (Fig. 1C), suggesting that Mp1 relocalises to the cytoplasm upon association with Mp58.

Computational modelling of the Mp1 and Mp58 structures with IntFOLD-TS, ModFOLD9, and DeepUMQA-X indicated that Mp1 and Mp58 both predominantly form α-helices (Fig. S2). The computationally predicted structure of Mp1 (ModFOLD9: p = 0.001, E = 7.93e^-4^, GMǪS: 0.571, DeepUMQA-X: TM-score: 0.734 and Global-lDDT: 0.614) features a helix-turn-helix consisting of two interacting α-helices; a 34-residue helix at Thr23-Tyr56 (H1) and 40-residue helix at Thr63-Thr102 (H2). There are 47 residue pairs that interact between the two helices (H1: 20, H2: 18, all located internal to their respective helices) with a distance-of-closest approach equal to 7.4 Å. The predicted Mp58 structure (ModFOLD9: p = 0.01, E = 7.26e^-3^, GMǪS: 0.4532, DeepUMQA-X: TM-score: 0.569 and Global-lDDT: 0.566) consists of three α-helices and one 3_10_ helix; a 38-residue α-helix at Leu6-Phe43 (H1) which interacts with the 32 residue α-helix at Tyr49-Phe80 (H2) and the 3 residue 3_10_ helix at Met98-Asn100 (H3) that interacts with the 18 residue α-helix at Asp102-Met119 (H4). There are a total of 48 internally sited interacting residue pairs between H1 and H2 (21 and 19 residues involved respectively) with a distance-of-closest approach of 7.6 Å and a second helix-helix interaction between 7 residues pairs of H2 and H4 (5 residues from each helix) with distance of 12.8 Å.

To understand how Mp1 and Mp58 interact to form an effector complex, we performed further modelling of the proteins using AlphaFold3, MultiFOLD, and DeepUMQA-X multimer quality analysis. Our models produced potential Mp1-Mp58 complex combinations with accuracy scores which ranged from ipTM 0.19 - 0.71 for interface predicted template modelling (iPTM) and 0.41 - 0.71 for predicted template modelling (pTM) (Fig. S3). Analysis of the Mp1-Mp58 complex with a 1:2, 2:1, 2:2, and 4:4 ratio produced models with the highest confidence (Fig. 1D & S4), indicating that Mp1 and Mp58 may form a larger oligomeric complex.

The Mp1-Mp58 modelled interaction with 1:2 stoichiometry identifies a more involved interaction between Mp58-Mp58 chains involving a total of 89 interface residues with an interface area of 2552 Å^2^ and 2536 Å^2^ for each respective chain, in contrast to the Mp1-Mp58 interactions with chains B and C which involve a sum of 24 and 33 interface residues and a ≈26– 38% chain-dependant interaction surface area. In this model, both Mp58 H1 helices are aligned anti-parallel and this is the case for H2-H2 and H3-H3 helices as well. The Mp1 H1 α-helix interacts with the Mp58(H1)-Mp58(H1) α-helix bundle at ≈45° forming 5 salt bridges and 14 hydrogen bonds (Fig. S5A). A similar preference for Mp1-Mp1 interaction can be seen in the 2:1 stoichiometric model, where there are a total of 88 interface residues with an interaction surface area of 2199 Å^2^ and 2207 Å^2^ for each compared to the two Mp1-Mp58 interactions with 22 and 21 interface residues with a ≈29% the interaction surface area. Each Mp1 α-helix aligns anti-parallel to the corresponding same α-helix in the interacting Mp1 chain (H1-H1, H2-H2) while Mp58 H1 interacts with the Mp1-Mp1, H1-H1 helical bundle at an angle of ≈45° with 3 salt bridges and 8 hydrogen bonds (Fig. S5B). The 2:2 heterodimer exhibits the same features where Mp1 and Mp58 interact more with other copies of the same protein, with each α-helix interacting with its counterpart in the opposing molecule in an anti-parallel fashion. The Mp1 H1-H1 helical pair interact with the Mp58 helical pair in the same manner as before at an ≈45° angle (Fig. S5C). The 4:4 heterodimer interaction builds upon the 2:2 structure by the addition of a second 2:2 structure that is spatially antisymmetric (Fig. S5D). Of note, Mp58-Mp58 H2-H2 interactions do not occur with the adjacent H2 but rather with the diagonally opposed H2 helix.

### Mp1 and Mp58 transient co-expression weakly enhances host susceptibility

Previous experiments to implicate Mp1 and Mp58 effector activity in plant-aphid interactions have been based on ectopic expression in either Arabidopsis or *N. benthamiana* followed by measurements of aphid performance (Elzinga et al., 2014; Pitino and Hogenhout, 2013; Rodriguez et al., 2017). Given the evidence Mp1 and Mp58 interact with one another, we investigated whether the combined expression of the effector pair alters host susceptibility to aphids. We previously showed that Mp1 ectopic expression in *N. benthamiana* enhanced susceptibility only when Mp1 was expressed under a phloem-specific AtSUC2promoter (Rodriquez et al.., 2017) while ectopic expression of Mp58 reduced aphid performance in transgenic Arabidopsis lines when expressed under a 35S and AtSUC2 promoter (Elzinga et al., 2014), and under a 35S promoter in *N. benthamiana* (Escudero-Martinez et al., 2018). Therefore, we opted to use the phloem-specific AtSUC2 promoter to express Mp1 and Mp58 both individually and together, in *N. benthamiana* and challenged infiltrated leaf areas with *M. persicae* to determine the combined impact of these effectors on host susceptibility. In line with previous work, Mp1 expression resulted in an increase in fecundity compared to the vector control (Fig. 1E). Expression of Mp58 did not result in a significant decrease in aphid fecundity compared to the vector control. When Mp1 and Mp58 were co-expressed, we observed a trend towards increased fecundity, however this increase was not statistically significantly different to the vector control (Fig. 1E).

### The Mp1-like and Mp58-like interaction is conserved in cereal aphid *Rhopalosiphum padi*

With the genetic linkage and shared transcriptional control of the Mp1-Mp58 pair being conserved across different aphid species (Thorpe et al., 2018), we were interested to test whether Mp1-like and Mp58-like from other aphids can also interact. We co-expressed the putative Mp1/Mp58 orthologues from the aphid *Rhopalosiphum padi* (bird cherry-oat aphid) in *N. benthamiana* and performed co-IP to detect a possible interaction. We were able to detect GFP-Rp58 upon immunoprecipitation of myc-Rp1 but not in the myc-GUS negative control, indicating the interaction of the effector pair is conserved across aphid species (Fig. 2A).

**Figure 2:**
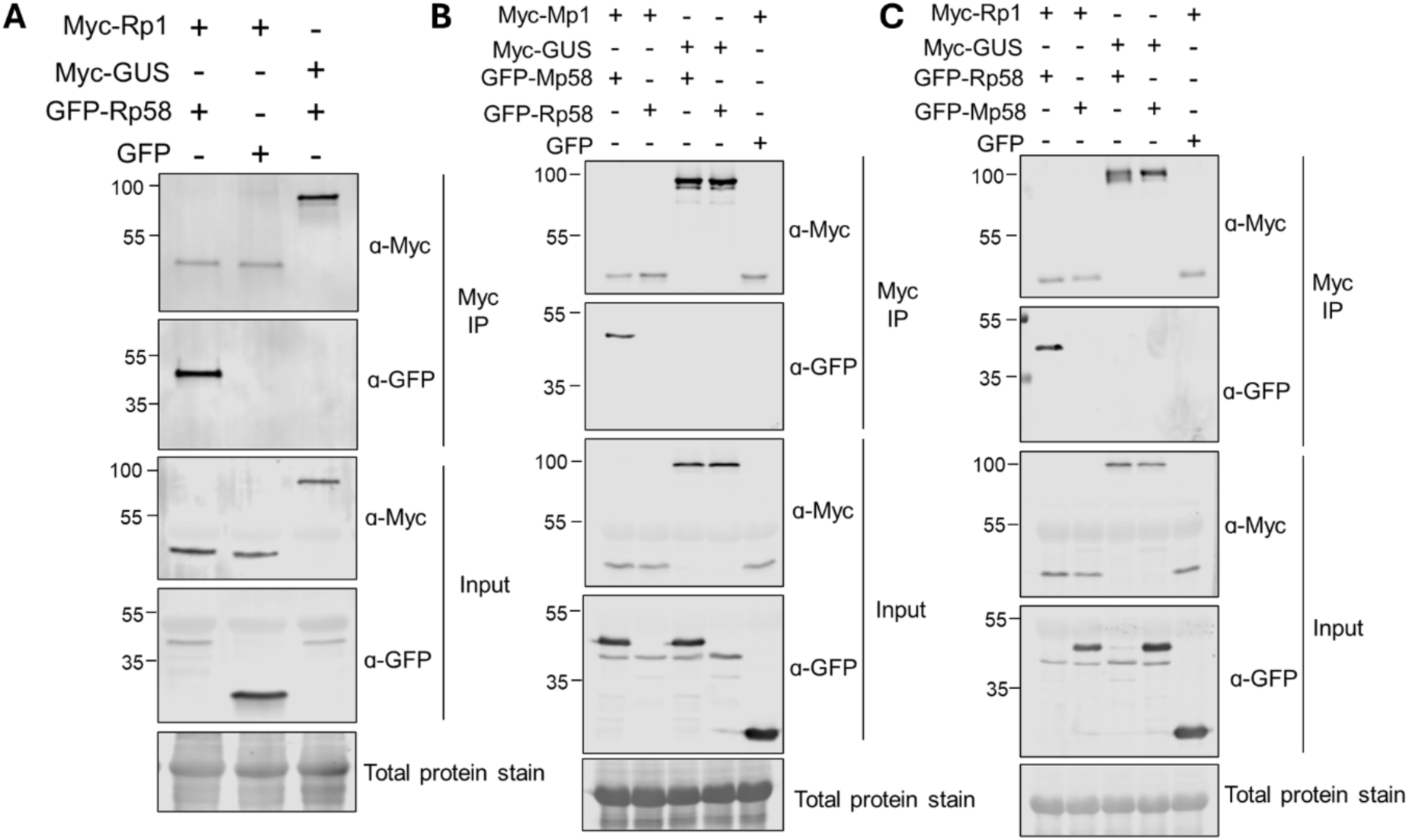
Mp1-like and Mp58-like interaction is conserved in the cereal aphid *Rhopalosiphum padi,* but interspecies interaction of the Mp1-like, Mp58-like effector pair is not detected. **(A)** Myc-Rp1 and GFP-Rp58 were transiently expressed via Agroinfiltration with each other or GFP and myc-GUS as controls. Proteins were pulled down via myc-trap **(B)** Myc-Mp1 and **(C)** Myc-Rp1 were co-expressed in *N. benthamiana* with GFP-Mp58/GFP-Rp58 or GFP and myc-GUS as controls. Proteins were pulled down via myc-trap. Ladders represent kDa.

With Rp1 and Mp1 showing 56% identity and Rp58 and Mp58 showing 65% identity (Escudero-Martinez et al., 2020), we asked the question as to whether complex formation can take place with pair members across aphid species. We co-expressed myc-Mp1 or myc-Rp1 with GFP-Mp58 or GFP-Rp58 and performed a co-IP to test for interaction. We were only able to detect GFP-Mp58 but not GFP-Rp58 upon immunoprecipitation of myc-Mp1 (Fig. 2B). Likewise, we co-expressed myc-Rp1 with GFP-Rp58 or GFP-Mp58 and only detected GFP-Rp58 but not GFP-Mp58 upon immunoprecipitation of myc-Rp1 (Fig. 2C). Our observations point to species specificity in Mp1(-like) and Mp58(-like) complex formation, suggesting that the effectors have evolved together in each aphid species to mediate their activities.

### Mp1 and Mp58 are both able to associate with host protein VPS52

We previously showed that Mp1 associates with VPS52 from Arabidopsis (AtVPS52) and *S. tuberosum* (StVPS52) in a species-specific manner and that this association is linked to Mp1 virulence activity (Rodriguez et al., 2017). Since we found that Mp1 forms a complex with Mp58, and Mp1 and Mp58 share similar secondary structural features, we considered the possibility that Mp1 and Mp58 may share targets. We therefore tested whether Mp58, like Mp1, can associate with AtVPS52, and whether a possible association would be dependent on the presence of Mp1. GFP-Mp58 and RFP-AtVPS52 were co-expressed in *N. benthamiana* in the presence or absence of myc-Mp1 by agroinfiltration, followed by immunoprecipitation of RFP-AtVPS52. Both in the presence and absence of myc-Mp1, we were able to detect GFP-Mp58 upon immunoprecipitation of RFP-AtVPS52, indicating that Mp58 indeed associates with AtVPS52, and that this association is independent of Mp1 (Fig. 3A). The level of GFP-Mp58 that was detectable upon RFP-AtVPS52 pull-down was similar in the presence versus absence of myc-Mp1, suggesting that the presence of Mp1 does not strengthen/weaken the association of Mp58 with AtVPS52. Likewise, we tested whether the interaction between Mp1 and AtVPS52 is altered in the presence of Mp58. We co-expressed myc-Mp1 and RFP-AtVPS52 in the presence or absence of GFP-Mp58. As expected, and in line with our previous work, myc-Mp1 was detected upon immunoprecipitation of RFP-VPS52 (Fig. 3B). The presence of GFP-Mp58 did not visibly affect the level of detectable myc-Mp1 in the RFP-VPS52 pull-down. Overall, these observations indicate that both Mp1 and Mp58 can interact independently with AtVPS52.

**Figure 3:**
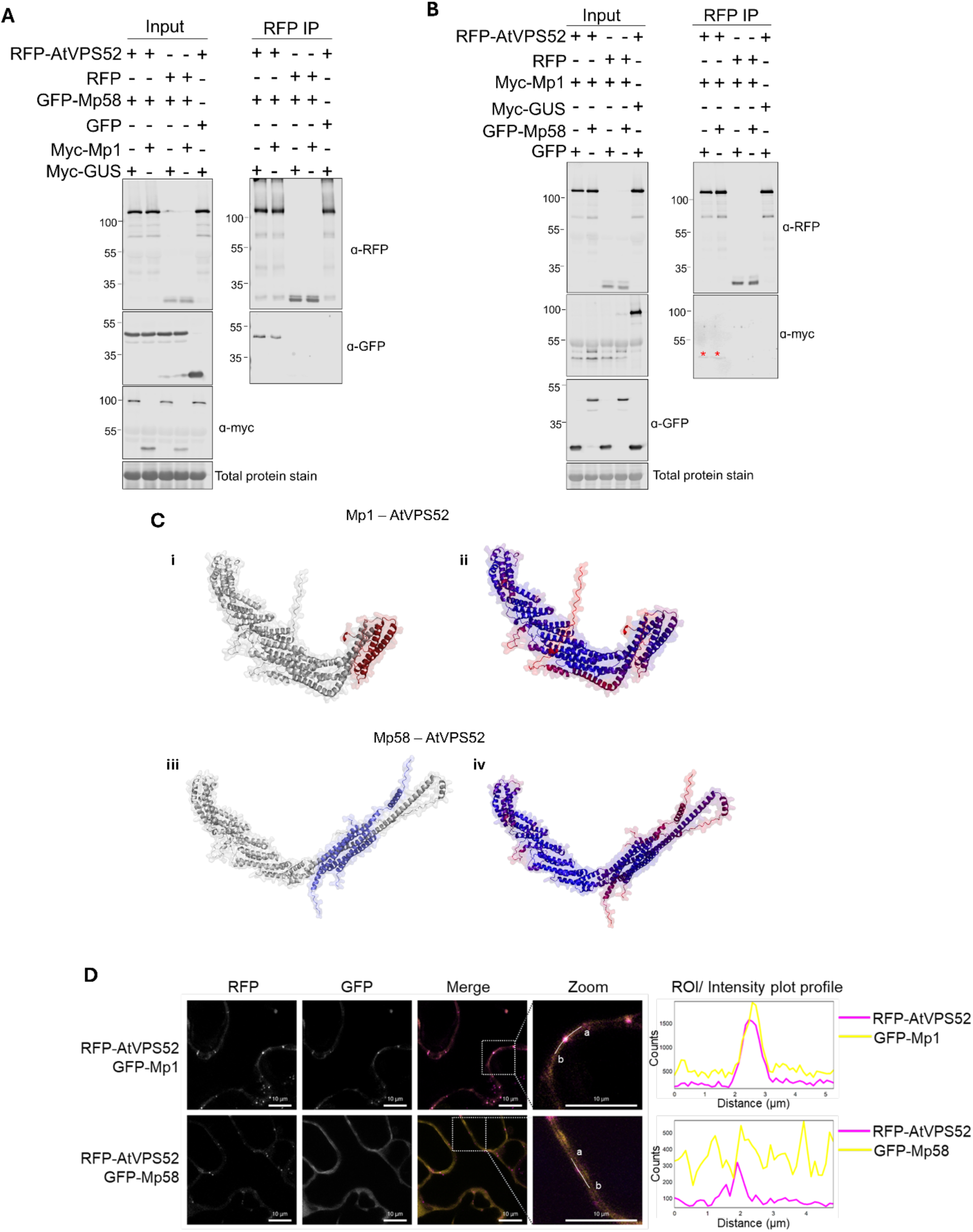
Mp1 and Mp58 both independently interact with AtVPS52, but only Mp1 localises to AtVPS52-associated vesicles. Immunoprecipitation of RFP-AtVPS52 with **(A)** GFP-Mp58 in the presence of myc-Mp1 or myc-GUS (control) or **(B)** myc-Mp1 in the presence of GFP-Mp58 or GFP (control). Proteins were transiently expressed in *N. benthamiana* via agroinfiltration and pulled down with RFP-trap. Ladders represent kDa **(C)** AlphaFold3 multimeric prediction of Mp1-AtVPS52 (**i.** and **ii.**) with 1:1 stoichiometry with AF3-derived ipTM: 0.52, pTM: 0.65 and DeepUMQA-X derived TM-score: 0.816, QS-score: 0.701, Global-lDDT:0.665 and Interface-lDDT: 0.632. Figure **i.** shows the Nʹ terminal of VPS52 (grey) interacting with Mp1 (red) and **ii.** shows the same interaction coloured by plDDT on a red-blue / low-high confidence scale. Figures **iii.** and **iv.** show AlphaFold3 multimeric prediction for Mp58-AtVPS52 with a 2:1 stoichiometry, with an AF3-derived ipTM: 0.20, pTM: 0.52 and DeepUMQA-X derived TM-score: 0.748, QS-score: 0.371, Global-lDDT:0.677 and Interface-lDDT: 0.642. Figure **iii.** shows a potential interaction between the Nʹ terminal of VPS52 (grey) and Mp58 (blue) and **iv.** Shows the interaction coloured by plDDT. **(D)** Subcellular localisation of RFP-AtVPS52 (in magenta) with GFP-Mp1 or GFP-Mp58 (in yellow). Proteins were transiently expressed *N. benthamiana* via agroinfiltration and localisation was observed with confocal microscopy. Magnification of 60X (water immersion lens), scale bar = 10 µm (controls of individually expressed proteins Fig. S8). Presented images are single plane images. The merged panel transect correspond to line intensity plot showing fluorescence distribution across the marked locus.

Predicted structural modelling of the Mp1-AtVPS52 interaction (AF3: ipTM = 0.52 and pTM = 0.65. DeepUMQA-X: TM-score: 0.816, QS-score: 0.701, Global-lDDT:0.665, Interface-lDDT: 0.632) results in an interaction between the Nʹ-terminal portion of AtVPS52 and the helical bundle of Mp1 with a total of 76 interface residues, 2 salt bridges and 6 hydrogen bonds with a total interface area of 2283 Å^2^ of VPS52 and 2145 Å^2^ of Mp1. The interaction elicits a break in the α-helix of the Nʹ-terminal portion of VPS52 between residues aa128-134, resulting in an H4 α-helix (E60-E128), β-turn at I129-I132 (IGSI), H5 3’10-helix (S134-I136) and H6 α-helix (L137-I167) of which the portion beyond the Cʹ-terminus of the helix interacts with residues internal to H4 with a total of 34 interacting residues at a distance of 6.1 Å. The VPS52-H4 helix interacts with Mp1-H1 (22 residues, distance 1.9 Å) and Mp1-H2 (4 residues, distance 10.7 Å) while VPS52-H6 predominantly interacts with Mp1-H2 (18 residues, distance 11.0 Å) then secondarily with Mp1-H1 (8 residues, 11.9 Å) (Figs. 3C, i & ii, S6A & S7A).

Modelling of the Mp58-AtVPS52 interaction (AF3: ipTM = 0.20 and pTM = 0.52. DeepUMQA-X: TM-score: 0.748, QS-score: 0.371, Global-lDDT:0.677, Interface-lDDT: 0.642) produced the best iPTM and TM scores with a 2:1 stoichiometry. This interaction does not disrupt the N-terminal H4 α-helix, which remains a single helix spanning E60-I167. The Mp58-Mp58 complex retains the same structure as previously described and this interacts with VPS52 residues L9-D214, with the main interaction occurring between R154-D214 with a total of 3 salt bridges and 8 hydrogen bonds, forming a AtVPS52 interface area of 3046 Å^2^ and 3080 Å^2^ with the Mp58-Mp58 complex (Figs. 3C, iii & iv, Fig. S6B & S7B).

### In contrast to Mp1, Mp58 does not re-localise to AtVPS52-associated vesicles

In plants, VPS52 has been found to localise to the Golgi and pre-vacuolar compartments (Guermonprez et al., 2008; Lobstein et al., 2004; Rodriguez et al., 2017). In our previous work, we demonstrated that in the presence of AtVPS52 or StVPS52, Mp1 relocalises to VPS52-associated vesicles (Rodriguez et al., 2017). As we show that Mp58, similar to Mp1, interacts with AtVPS52 (Fig. 3A) we assessed whether Mp58 and AtVPS52 also co-localise. We co-expressed GFP-Mp58 or GFP-Mp1 (positive control) with RFP-AtVPS52 individually to determine subcellular location. As expected, GFP-Mp1 localised to the nucleus and cytoplasm, but when expressed alongside RFP-AtVPS52, re-localised to AtVPS52-associated vesicles (Figs. 3D & S8). Interestingly, GFP-Mp58 localised to the nucleus and cytoplasm regardless of the presence of RFP-AtVPS52 (Fig. 3D & S8), indicating that Mp58, unlike Mp1, does not relocalise to VPS52-associated vesicles. These observations suggest that although Mp1 and Mp58 both interact with AtVPS52, only the Mp1-AtVPS52 interaction occurs at VPS52-associated vesicles when we co-express with one effector at a time.

### Mp1 and Mp58 interact with different regions of AtVPS52

Based on our predicted 3D protein structure models (Fig 3C), we generated AtVPS52 constructs expressing the 170 N-terminal amino acids of the protein, which contains the predicted interaction site for Mp1 (AtVPS52_1-170_, Fig. 4A), as well as a set of VPS52 chimeras in which we swapped this N-terminal domain between AtVPS52 and HvVPS52. With the Mp58-AtVPS52 interaction site less defined by computational modelling, we focused our structure-function analyses on AtVPS52 in the context of the Mp1 interaction whilst including Mp58 in various interaction assays. We first assessed interaction of AtVPS52_1-170_ with effectors Mp1 and Mp58 *in vitro*. For this, we individually expressed His-AtVPS52_1-170_ and GST-Mp1 or His-Mp58 and GST-AtVPS52_1-170_ in *E. coli*and combined lysates before IMAC. Purification of His-AtVPS52_1-170_ resulted in the co-purification of GST-Mp1 (Fig. 4B). We were unable to consistently purify GST-AtVPS52_1-170_ with His-Mp58, suggesting that the Mp58-AtVPS52 features different structural requirements. Multiple attempts to detect the tagged the AtVPS52_1-170_ protein domain when transiently expressed in *N. benthamiana* were unsuccessful, possibly due to low protein stability in plant cells. Therefore, to test for interaction of the N-terminal AtVPS52 domain with the effectors, we designed VPS52 chimeras where the N-terminal region of AtVPS52 was swapped with the N-terminal region of HvVPS52, which is unable to interact with Mp1 (Rodriguez et al., 2017), and vice versa (Fig. S9). We refer to these chimeras as HvVPS52^At^ ^1-170^ and AtVPS52^Hv1-165^, respectively (Fig. 4A). GFP-Mp1 or GFP-Mp58 were co-expressed with RFP-tagged full-length AtVPS52, HvVPS52, or the VPS52 chimeras in *N. benthamiana* followed by immunoprecipitation of RFP-VPS52 variants. As expected, GFP-Mp1 co-immunoprecipitated with RFP-AtVPS52 but not HvVPS52 (Fig. 4C). GFP-Mp1 also co-immunoprecipitated with RFP-HvVPS52^At1-170^, but not with RFP-AtVPS52^Hv1-165^ (Fig. 4C). Although GFP-Mp58 co-immunoprecipitated with RFP-AtVPS52 as expected (Fig. 4D), we either observed no or a weak interaction of GFP-Mp58 with RFP-HvVPS52 across repeated experiments (Fig. 4D, S10), pointing to a potentially weak interaction between Mp58 and HvVPS52. Interestingly, GFP-Mp58 consistently co-immunoprecipitated with RFP-AtVPS52^Hv1-165^, while we observed either no interaction or a weak interaction with RFP-HvVPS52^At1-170^ across biological replicates (Fig. 4D, S10). Our observations indicate that while the Mp1-VPS52 interaction is host-specific and N-terminus dependent, the Mp58-VPS52 interaction may be more undiscriminating.

**Figure 4:**
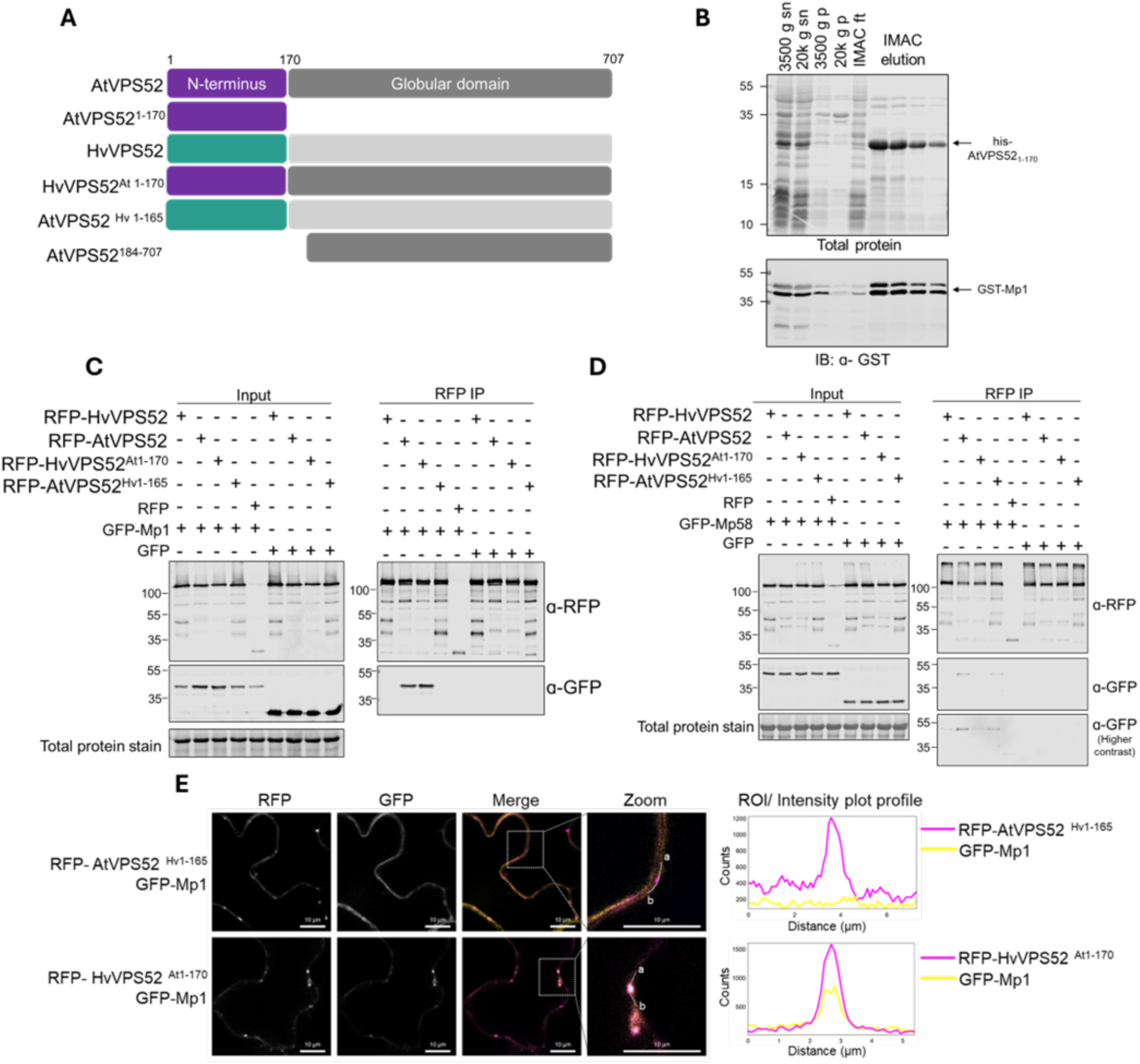
Mp1 and Mp58 interact with different regions of AtVPS52. **(A)** Schematic of full-length Arabidopsis (At) and barley (Hv) VPS52, and VPS52 variants **(B)** co-purification of His-AtVPS52 1-170 with GST-Mp1 from recombinant protein expressed in *E. coli*. Expression of His- and GST-tagged proteins were induced in *E.coli* individually and protein lysates were combined. Protein was purified via IMAC and blotted against GST. Sn - supernatant; p – pellet; ft – flowthrough; IMAC – immobilised metal affinity chromatography. **(C)** and **(D)** GFP-Mp1 or GFP-Mp58 were co-expressed with RFP-HvVPS52, RFP-AtVPS52, or N-terminal chimeras in *N. benthamiana* via agroinfiltration. RFP-tagged proteins were pulled-down with RFP-trap and blotted against GFP. RFP and GFP were used as negative controls. Ladders represent kDa **(E)** GFP-Mp1 (in yellow) was co-expressed with RFP-AtVPS52^Hv1-165^ or RFP-HvVPS52^At1-170^ (in magenta) in *N. benthamiana* via agroinfiltration and subcellular localisation and localisation was observed with confocal microscopy. Magnification = 60X (water immersion lens), scale bar = 10 µm (controls of individually expressed proteins Fig. S11). Presented images are single plane images. The merged panel transect correspond to line intensity plot showing fluorescence distribution across the marked locus.

To determine whether the VPS52 chimeras feature a similar subcellular localisation to wild-type AtVPS52, we performed confocal microscopy and found that both RFP-HvVPS52^At1-170^and RFP-AtVPS52^Hv1-165^ localised to vesicles (Fig. S11). Since only Mp1 and not Mp58 re-localised to wild-type AtVPS52-associated vesicles, we performed co-localisation experiments of the VPS52 chimeras with Mp1 only. In line with previous observations (Rodriguez et al., 2017), GFP-Mp1 localised to RFP-AtVPS52, but not RFP-HvVPS52-associated vesicles (Fig. 4E). When GFP-Mp1 was co-expressed with the RFP-VPS52 chimeras, we observed localisation of GFP-Mp1 to RFP-HvVPS52^At1-170^, but not RFP-AtVPS52^Hv1-165^-associated vesicles (Fig. 4E), suggesting that Mp1 association with AtVPS52 indeed requires the AtVPS52 N-terminal region.

We also generated a construct that expresses AtVPS52 missing the 170 N-terminal amino acids (AtVPS52^184-707^) for co-expression experiments. However, RFP-AtVPS52_184-707_ did not localise to vesicles (Fig. S12), suggesting that the N-terminal region of AtVPS52 is required for VPS52 association with vesicles and potentially its function in plant cellular trafficking. Overall, our data demonstrates that the N-terminal region of AtVPS52 is required for association with GFP-Mp1, highlighting that sequence variation within this region defines interaction specificity.

### An Mp1-Mp58-VPS52 multiprotein complex associates with vesicle-like structures

Although Mp1 and Mp58 can interact with AtVPS52 independently of each other, we show that these proteins also form an effector complex, raising the question as to whether the two effectors can associate with VPS52 at the same time to form a multi-effector-VPS52 complex.

We hypothesised if indeed all three interact together, we would be able to detect the different proteins when co-expressed in plant cells in the same subcellular vesicle-like structure observed for AtVPS52. We transiently expressed GFP-AtVPS52 with BFP2-Mp58 and/or RFP-Mp1 (with GFP, BFP2, or RFP as controls) in *N. benthamiana* and observed subcellular localisation using confocal microscopy. As previously shown, when AtVPS52 is expressed alone, we observed localisation to vesicle-like structures (Fig. 5). In line with our previous findings, we observed localisation of RFP-Mp1 to GFP-AtVPS52 associated vesicle-like structures, while BFP2-Mp58 remained localised to the nucleus and cytoplasm when co-expressed with GFP-AtVPS52 (Fig. 5). Interestingly, when GFP-AtVPS52, BFP2-Mp58, and RFP-Mp1, were co-expressed we observed partial localisation of Mp1, Mp58, and AtVPS52 to AtVPS52-associated vesicle-like structures (Fig. 5). This suggests that Mp1, Mp58, and AtVPS52 may indeed form a complex and that Mp58 only localises to AtVPS52-associated vesicles in the presence of Mp1, pointing to potential effector cooperation.

**Figure 5:**
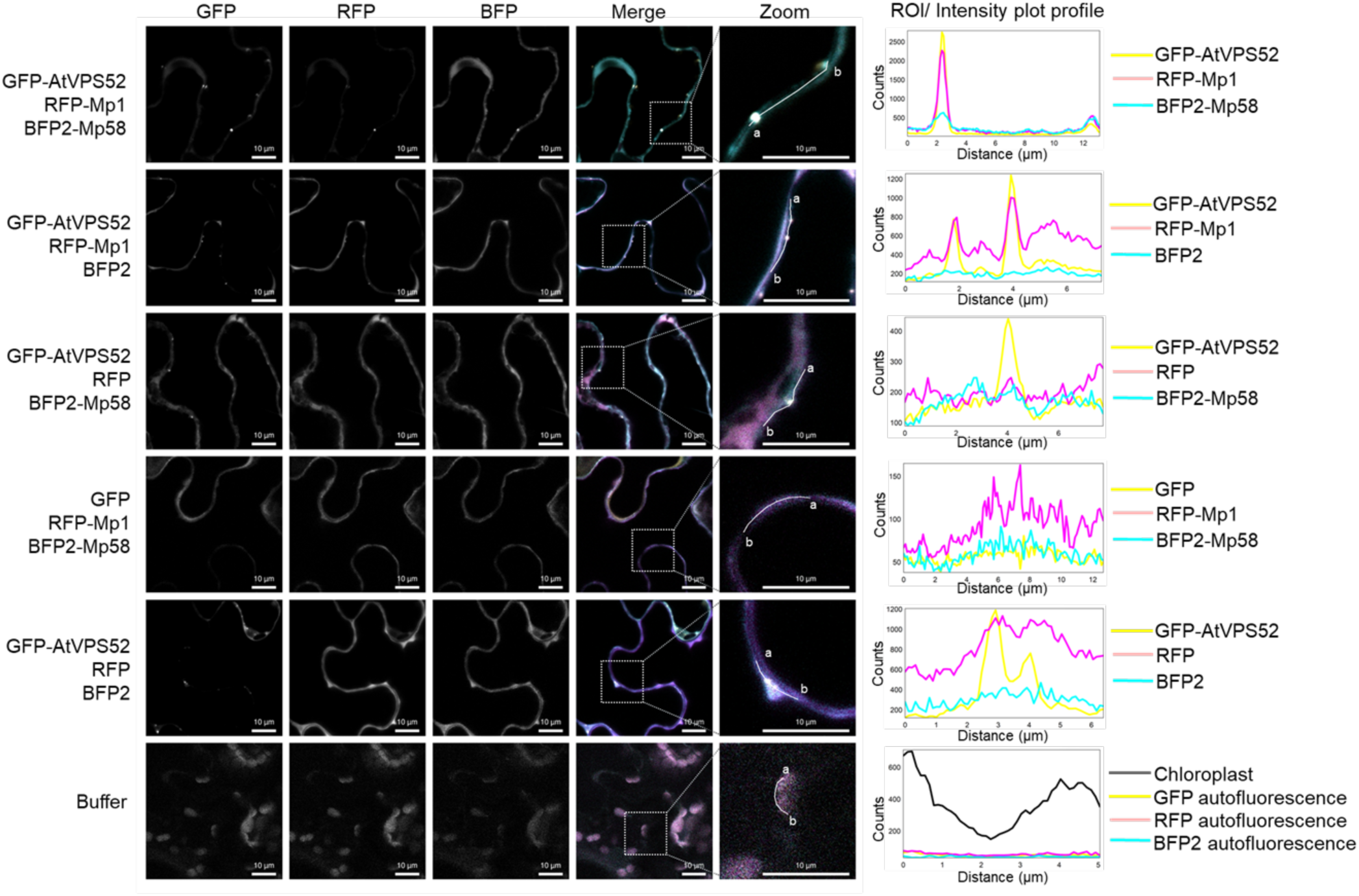
AtVPS52, Mp1, and Mp58 co-localise to vesicle-like structures. GFP-AtVPS52 (in yellow) was co-expressed with RFP-Mp1 (in magenta) and/or BFP2-Mp58 (in cyan) (GFP, RFP, and BFP2 used as controls). Proteins were expressed transiently in *Nicotiana benthamiana* and imaged 2dpi. Magnification = 60X (water immersion lens), scale bar = 10 µm. Presented images are single plane images. The merged panel transect correspond to line intensity plot showing fluorescence distribution across the marked locus.

## Discussion

Effectors play a crucial role in manipulating plant cell processes to facilitate pathogen infection and pest infestation. While pathogens and pests typically secrete a wide range of effectors into their host plants, studies on effector functions usually focus on individual proteins. Here, we demonstrate that effectors from the economically important aphid pest, *M. persicae*, form a complex to potentially work together, underscoring the importance of examining effectors within the context of the wider effector repertoire. We show that effectors Mp1 and Mp58, which are encoded by a physically linked and tightly co-regulated gene pair conserved across aphid genomes, interact to form an effector complex. Together these effectors can bind to the previously identified Mp1 target VPS52 at vesicle-like structure (Rodriguez et al., 2017). Whilst effector complexes were reported in plant pathogenic fungi (Cao et al., 2018; Ludwig et al., 2021; Yu et al., 2024) their biological function and relevance are largely unknown. Our findings point to effector complex formation in plant-insect interactions and highlight a yet-to-be-explored layer of complexity in the plant-insect molecular dialogue.

Effector complex formation has been reported in pathogens more broadly and was first described for effectors LubX and SidH in the human bacterial pathogen *Legionella pneumophila*. Effector LubX, an ubiquitin E3 ligase, targets the SidH effector for proteasome-mediated degradation to regulate its activity during infection stages (Kubori et al., 2010). The term ‘metaeffector’ was proposed to describe effectors like LubX that regulate the activities of other effectors (Joseph and Shames, 2021; Kubori et al., 2010). Extensive interaction analyses of ∼330-390 *L. pneumophila* effectors have since identified ∼20 interacting pairs pointing to extensive metaeffector activity (O’Connor Mount et al., 2024; Urbanus et al., 2016). Similar, some effectors from plant pathogenic microbes feature metaeffector activity, for example to suppress activation of ETI (Martel et al., 2022). This can be achieved via protein-protein interactions with the same host protein, as is the case for several *Pseudomonas syringae* effectors (reviewed in Bundalovic-Torma et al. (2022)). Further evidence for metaeffector activity in plant-pathogen interactions is provided by evidence that some effectors can form heteromeric complexes, which are suggested to be involved in either regulation of effector translocation and/or activity (Alcântara et al., 2019; Cao et al., 2018; Li et al., 2016; Ludwig et al., 2021; Ma et al., 2015). Similar to the Mp1-Mp58 effector pair in *M. persicae*, interacting and/or cooperating effectors *from Fusarium oxysporum f. sp. Lycopersici* (Fol) and *f. sp conglutinans* are co-located, however, their expression seems to be divergent rather than co-regulated (Ayukawa et al., 2021; Ma et al., 2015; Yu et al., 2024). Functional studies showed that *Fol* effector Six5 interacts with Avr2, and increases plasmodesmata (PD) size exclusion limit to facilitate the symplastic transport of Avr2, and potentially additional effectors, a function that is conserved amongst plant fungal pathogens (Blekemolen et al., 2022; Cao et al., 2018; Talbi et al., 2024, 2023). While we demonstrate that Mp1 and Mp58 physically interact with one another as well as with the previously identified Mp1 target VPS52, the mechanistic role and composition of effector(-target) complex(es) remains to be addressed. Our observations raise new questions regarding when and where complex formation takes place, before, during, and/or after effector delivery (Fig. 6), and what the biological relevance of complex formation is. For example, Mp1-Mp58 complex formation may regulate individual effector activities towards certain host targets, regulate effector delivery, and/or may allow diversification of potential host targets (i.e. complexes may interact with different host proteins than individual effectors). The individual Mp1 and Mp58 effector proteins are predicted to be relatively unstructured with two regions predicted to form α-helices, though these predictions have relatively low accuracy scores (Waksman et al., 2024). Assuming a 1:1 interaction, computational modelling highlights an Mp1-Mp58 interaction interface between the predicted α-helices. However, our computational analyses indicate that Mp1 and Mp58 likely form a larger oligomeric complex.

**Figure 6:**
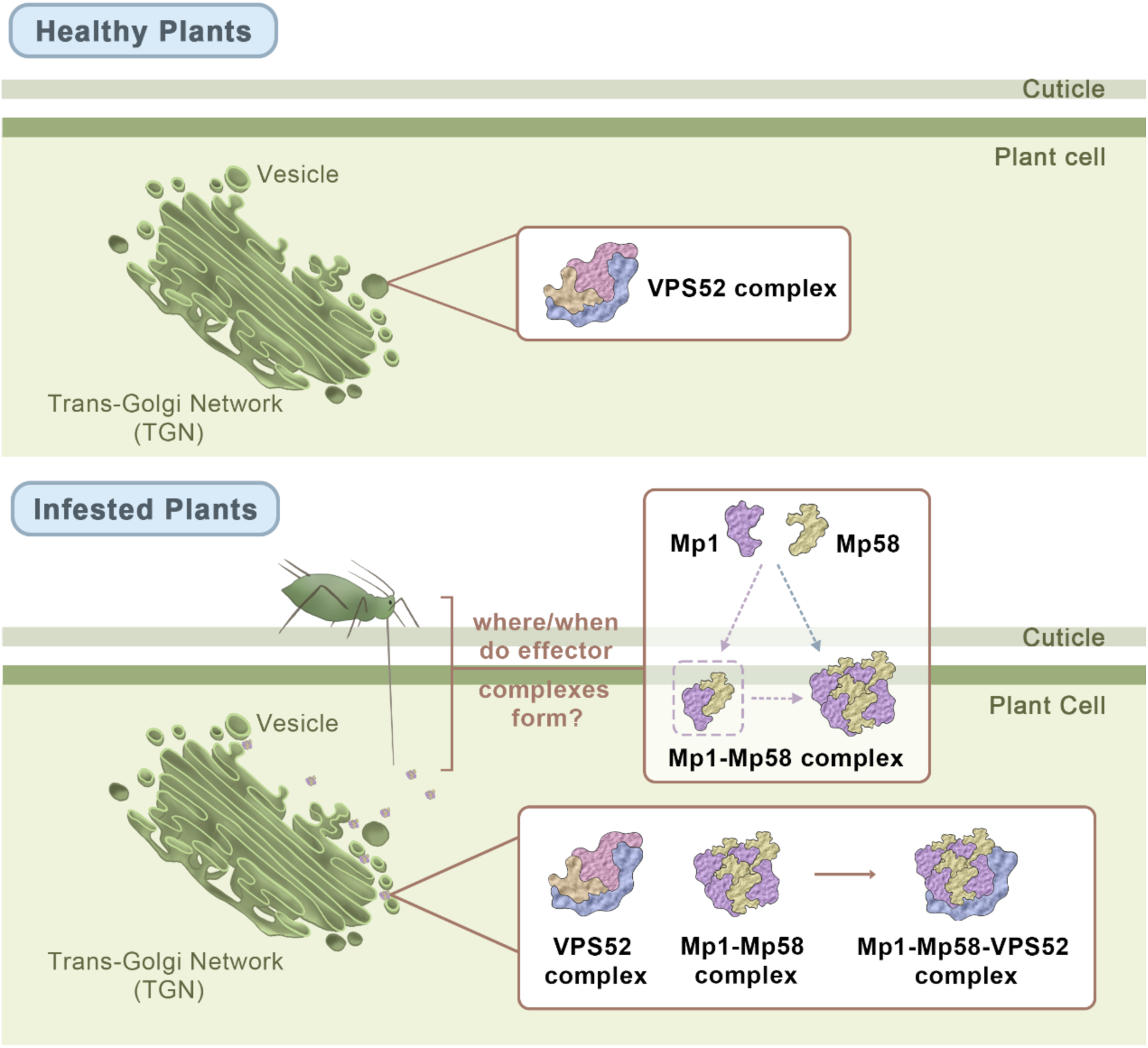
Working model of Mp1-Mp58-AtVPS52 complex. Upon infestation, *Myzus persicae* effectors, Mp1 and Mp58, are secreted into host plant cells. Mp1 and Mp58 interact to form a complex, but the size and stoichiometry of the complex and whether the complex forms before, during, or after delivery remains to be investigated. Mp1 and Mp58 associate with VPS52-associated vesicles, which may disrupt or alter VPS52-associated complexes to alter vesicle trafficking mechanisms to benefit aphid feeding.

Although the Mp1-Mp58 gene pair is co-located and conserved across the genomes of at least 5 aphid species, with evidence for shared transcriptional control, our data show that the effector-effector interaction is species-specific. The sequence variation of the Mp1 and Mp58 effector sequences across species (±55-65% sequence identity between the *M. persicae* and *R. padi* pair) suggests the effector pairs have evolved together within each aphid species and diversified across species due to functional adaptation in different host plants species. Either the host protein(s) targeted by the effector pair in different host species have a similar function but feature significant structural diversity or the effector pair may have evolved divergent functions in different hosts. Our previous work showed that Mp1 specifically targets VPS52 in host plant species, however we were unable to detect interaction between Rp1 and the HvVPS52 (Rodriguez et al., 2017), which may suggest that Mp1-like and Mp58-like effector pairs have evolved different functions across plant species, though further experimental work will be required to explore this.

The co-localisation of Mp1, Mp58, and AtVPS52 in vesicle-like structures upon co-expression is in line with a model that the two effectors and their target can associate on one complex (Fig. 6). Although both effectors also individually can associate with AtVPS52, the presence of Mp58 at AtVPS52-associated vesicle-like structures only occurs in the presence of Mp1. This raises the question whether Mp1 may be acting as a chaperone to Mp58 and/or facilitating its activity. In the case of the *Fol* Avr2-Six5 pair, the movement of Avr2 between cells is significantly increased in the presence of Six5, showing an important role of Six5 is to facilitate Avr2 movement and activity (Blekemolen et al., 2022). Whether, and if so, how Mp1 or Mp58 similarly regulate movement and/or activity of the other effector pair member remains to be investigated.

Additionally, Mp1 and Mp58 interact with different regions of AtVPS52, indicating that these effectors may simultaneously target the same host protein by binding to distinct domains (Fig. 4). We did not find evidence for competitive binding of Mp1 and Mp58 to VPS52 in Co-IP assays, which is in line with these effectors binding to different AtVPS52 regions. Although we previously showed the VPS52 over-expression increases host immunity to aphids, the mechanisms by which Mp1 and Mp58 target VPS52 and the downstream consequences of this remain to be revealed. Within the GARP complex VPS52 associates with the subunits VPS51, VPS53 and VPS54, and in *Saccharomyces cerevisiae* these interactions involve the N-terminal α-helices of these proteins, as determined by electron microscopy (Chou et al., 2016). However, most of these subunits, including VPS52, can also associate with additional cellular trafficking complexes, such as the EARP (endosome-associated recycling protein) complex which is involved endocytic recycling as shown in human cells and yeast (Schindler et al., 2015). We hypothesise that Mp1 and/or Mp58 may disrupt or modify one of these cellular trafficking complexes through association with VPS52 to benefit aphid infestation (Fig. 6). In line with this, we previously found that AtVPS52 was degraded post-transcriptionally upon aphid infestation, however, this observation could not be attributed to Mp1 based on transient ectopic expression-based assays (Rodriguez et al., 2017). Further biochemistry approaches will be needed to reveal whether and how Mp1/Mp58 may affect cellular trafficking pathways via VPS52, and if so, how this involves effector cooperation.

The lack of a strong virulence phenotype upon combined expression of Mp1 and Mp58 may point to a role of effector complex formation in perhaps effector activity regulation rather than cooperation. However, the functional assays available to test whether *M. persicae* effectors contribute to promoting susceptibility have significant limitations as they rely on ectopic expression of aphid proteins in plants. It is well established that ectopic expression of effectors in plants may result in excessive or mis-targeting of host proteins, which can lead to different or even opposite phenotypes to those expected based on actual virulence activity, this in part to a lack of knowledge on endogenous effector amounts and spatiotemporal effects. Moreover, our combined effector expression assay requires co-transformation of the same plant cells using *Agrobacterium* carrying different constructs, which may dilute any observable phenotypes. In previous studies (Pitino and Hogenhout, 2012; Rodriguez et al., 2017), expression of Mp1 alone increased host susceptibility to aphids, whereas expression of Mp58 reduced host susceptibility. Differences in experimental conditions, such as the expression of Mp58 under the AtSUC2 promoter in *N. benthamiana*, which has not been previously tested, could explain the lack of an Mp58 phenotype in our assays. With no genetic modification system currently established for *M. persicae* and RNAi resulting in low or inconsistent levels of reduced expression, options to reliably screen for aphid effector virulence activities are limited with further tool development needed to help advance the field.

In conclusion, this work provides evidence that effectors from phloem-feeding insects can associate with one another to form effector complexes as well as with (shared) host proteins, potentially to promote infestation. Our data point to effector complex formation in plant-insect interactions and raises new research questions about the role of effector complex formation in regulating and/or diversifying effector activities, and as to whether effector complex formation is a common feature within herbivorous insect effector repertoires. Overall observation that a conserved and genetically linked effector pair can form an effector complex highlights a yet-to-be explored layer of complexity to consider in the molecular dialogue between plants and herbivorous insects.

## Materials and Methods

### Plant growth conditions and aphid maintenance

*Nicotiana benthamiana* plants were grown in a glasshouse with 16 h of light at ∼25°C.

*Myzus persicae* (Clone O) was maintained on *N. benthamiana* leaves in a growth cabinet with 16 hr light, 22°C during the day, 20 °C at night, 65% relative humidity.

### Plasmids

Constructs for *in planta* co-IP experiments and confocal assays had been cloned and described previously; GFP-Mp1 (pB7WGF2), RFP-AtVPS52, RFP-HvVPS52 (pK7WGR2) (Rodriguez et al., 2017), GFP-Mp58 (pB7WGF2) (Escudero-Martinez et al., 2020). Myc-Mp1 and myc-Rp1 (pGWB21) (Rodriguez et al., 2017). AtVPS52 _1-170_, AtVPS52 ^Hv1-165^, and HvVPS52 ^At1-170^ were synthesised by Twist Bioscience into the Gateway compatible vector pTwist ENTR. To generate RFP-tagged constructs for plant expression AtVPS52_1-170_, AtVPS52 ^Hv1-165^, and HvVPS52 ^At1-170^ were cloned into pK7WGR2 by LR reaction (LR clonase II, ThermoFisher 11791020) following the manufacturer’s instructions and transformed into *E. coli* Top10 cells. Constructs were verified by sequencing and transformed in *Agrobacterium tumefaciens* (GV3101) for expression in *N. benthamiana*. To generate BFP2-AtVPS52, previously cloned AtVPS52 in pEntr1A was cloned into pJRA109 (Allen et al., 2023) by LR reaction according to the manufacturer’s instructions and transformed into *E. coli* Top10 cells. Constructs were verified by sequencing and transformed in *Agrobacterium tumefaciens* (GV3101) for expression in *N. benthamiana*.

For aphid assays pAtSUC2::Mp1 (pHW59) had be cloned previously (Rodriguez et al., 2017). Mp58 was cloned from pEntr1A into pHW59 (Gottwald et al., 2000) by LR reaction to generate pAtSUC2::Mp58. Constructs were verified by sequencing and transformed into *Agrobacterium* (GV3101) for expression in *N. benthamiana*.

For *E. coli* protein expression and purification, codon-optimized GST, AtVPS52, Mp1 and Mp58 coding DNA sequences were synthesised by Integrated DNA Technologies or Twist Bioscience. Regions of interest were PCR-amplified and assembled into a pET-15b expression vector using GeneArt™ Gibson Assembly HiFi Master Mix (Thermo Fisher Scientific). 6xHis tags and TEV or HRV 3C protease cleavage sites were encoded in PCR primers.

### *Agrobacterium tumefaciens* infiltration for transient expression

*Agrobacterium tumefaciens* (strain GV3101) carrying the indicated constructs were grown shaking overnight at 28°C. Cultures were centrifuged at 2,500*g* for 10 mins at room temperature. Pellets were resuspended in infiltration buffer (10 mM MgCl_2_ and 10 mM MES, pH 5.6), the OD was adjusted (see below) and 200 µM acetosyringone was added to the infiltration buffer. For co-immunoprecipitation experiments Agrobacterium was diluted to OD_600_ 0.3 for all but GFP (OD_600_ 0.05) along with the silencing suppressor p19 (OD_600_ 0.1). For aphid performance assays the OD_600_ was 0.2 along with the silencing suppressor p19 (OD_600_ 0.1). For confocal-based co-localisation experiments, the OD_600_ ranged from 0.1-0.2 for all, along with the silencing suppressor p19 (OD_600_ 0.1 for Mp1-Mp58 co-localisation and 0.001 for all other experiments). Cultures were incubated for 2 hrs in the dark at 28°C. Three-four week-old *N. benthamiana* plants were infiltrated with Agrobacterium carrying the indicated constructs.

### Co-immunoprecipitation

Leaf tissue from *N. benthamiana* was ground in liquid nitrogen and was extracted with GTEN buffer (10% glycerol, 25 mM Tris-HCl pH 7.5, 1 mM EDTA) containing 0.2% Triton X-100, 10 mM DTT and 1x EDTA-free protease inhibitor cocktail (ThermoFisher; A32965). Samples were incubated for 10 minutes with frequent vortexing at 4°C. Samples were centrifuged at 13,000 rpm for 20 minutes. The supernatant was diluted with IP buffer (GTEN buffer, 0.1 % Tween-20) to reduce the DTT concentration to < 10 mM. lysate was incubated with either Chromtek myc-trap (Proteintech; ymta) or RFP-trap (Proteintech; rtma) rotating at 4°C for 2 hours. Beads were washed with IP buffer and protein was eluted at 80°C for 10 minutes in 2X SDS sample buffer containing 50 mM DTT. Samples were stored at −20°C until use.

### *E. coli* protein expression and purification

The pET-15b expression vector encoding protein of interest was transformed into *E. coli* BL21 (DE3) cells. Overnight culture was inoculated into 1 L lysogeny broth (LB) medium in a 2 L Erlenmeyer flask, and cell culture was grown at 37 °C until OD_600_ 0.4-0.8 was reached. The culture temperature was reduced to 25 °C, and protein expression induced via addition of isopropyl β-d-1-thiogalactopyranoside (IPTG) (final concentration 0.1 mM). After incubation at 25 °C for 4 hours, cells were harvested via centrifugation at 3500 g for 5 min, resuspended in ice-cold lysis buffer (25 mM HEPES, 150 mM NaCl, 25 mM imidazole, 1 mM DTT, pH 7.5) and stored at –20 °C. Thawed cell suspensions were lysed via sonication and subjected to centrifugation at 20,000 g for 30 min, and supernatant passed through 0.45 µm syringe filter. Proteins were purified from the solution via Ni^2+^ immobilized metal ion affinity chromatography, using Cytiva HisTrap FF column (Cytvia; 17525501) and ÄKTA go protein purification system.

### SDS-PAGE and immunoblot

Protein samples were incubated at 80°C for 10 minutes prior to loading on SDS-PAGE (10 or 15%) and run at 100-200V until the dye reached the bottom of the gel. Proteins were transferred to either nitrocellulose or PVDF membranes with a Tris-glycine transfer buffer (25 mM Tris, 192 mM glycine, 20% ethanol) for 90-120 mins at 90V or using the Bio-rad Turbo blot system. Membranes with input samples were stained for total protein prior to immunoblotting (see total protein staining). Membranes were blocked with 2% skimmed milk powder (Merck; 70166) in PBS-T (PBS containing 0.1 % Tween-20) before incubating with primary antibodies shaking 1-2 hrs at room temperature, or overnight at 4°C. The following antibodies were used: RFP (Chromotek; 5F8), GFP (Santa Cruz; sc-9996), myc (Santa Cruz; sc-40), GST (ThermoFisher; MA4-004). Membranes were washed with PBS-T before incubating with respective secondary antibodies, mouse (Licor; 926-33210) or rat (Licor; 926-32219) in the dark at room temperature for 1 hour. Proteins were detected with Licor Odyssey CLx in the 700 and 800 nm channels.

### Total protein staining

After transfer, membranes were dried for at room temperature (1 hour – overnight), rehydrated with ethanol (PVDF) or PBS (nitrocellulose) and stained with Revert™ 700 Total Protein Stain (Licor; 926-11021) or a homemade version (30% ethanol, 7% acetic acid, 0.001% Fast Green FCF [ThermoFisher; A16250]) for 10 minutes. Membranes were washed with Revert wash solution (30% ethanol, 7% acetic acid) to remove background and imaged using a Licor odyssey CLx with the 700 nm channel. Stain was removed with Revert stain stripper (30% ethanol, 100 mM sodium hydroxide). Membranes were rinsed with PBS before proceeding to blocking.

### Confocal imaging and image analysis

For Mp1 and Mp58 subcellular localisation experiments, confocal microscopy was performed 3 days post agroinfiltration. Live leaf tissue was mounted on a slide with the abaxial side facing upwards. Imaging was performed using a Zeiss LSM 710 with Plan Apochromat 40x/1.0 Water dipping lens. The excitation wavelength for mRFP was 561 nm, its emission was collected from 572 to 630 nm. GFP was imaged using 488 nm excitation, and its emission was collected from 500 to 530 nm. The pinhole was set to 1 AU for the shortest wavelength. Images were taken using line-by-line sequential scanning with 1024 × 1024 pixels with 2X line averaging.

For all other subcellular localisation experiments confocal microscopy was performed 2 days post agroinfiltration. For this, the abaxial side of the live leaf tissue was mounted on a 1% agar pad moulded on a slide using a gene frame sealed with a coverslip (22mm x 22mm #1.5 thickness (ThermoScientific™ - AB0577)). Imaging was performed using a Nikon A1R confocal microscope with CF1 Plan apochromat VC 60X water-immersion objective lens (NA-1.2, WD (0.27mm), coverslip correction set to 0.17 mm). The laser excitation for BFP2, GFP and RFP tags were at 405nm (violet diode laser), 488nm (argon laser-10mW), and at 561nm (sapphire laser-20mW), respectively. The emission ranges for BFP2, GFP and RFP were 425-475nm, 500 to 530nm, and 570 to 620nm, respectively. Autofluorescence from chlorophyll was captured at 488nm and emissions collected between 663 and 738nm. The pinhole was set to 1.2 AU for the shortest wavelength. Nyquist XY acquisition was performed for single optical slices, with zero offset and 5% laser power and a scanner zoom of 1.848 for all the channels. Galvano-line-by-line sequential scanning was employed with a pixel dwell time of 2.4 frames per second, and at 1024 × 1024 pixels with 2X line averaging. All the images were captured using NIS elements software provided by the Nikon. GFP fluorescence was shown in yellow to ensure accessibility for colour blind readers. All data were imported to our OMERO server, using OMERO. Insight (Version: 5.7.2) for organisation and annotation. Figures were generated in OMERO. Figure (Version: v4.4.3) (Allan et al., 2012), with brightness and contrast adjusted where necessary. ROI-based intensity profile plots were generated in ImageJ-Fiji (Version: 1.54g) using the plot profile plugin provided in the software.

### Protein structure and protein-protein interaction predictions

The protein sequences for Mp1 (XP_022169828.1) and Mp58 (XP_022169827.1) were submitted to SignalP 6.0, Phobius and DeepTMHMM for signal peptide and trans-membrane domain identification. By consensus a signal peptide was identified for Mp1 at aa19-20 with probability: 1.00 and cleavage probability: 0.99 resulting in a 120aa sequence and for Mp58 at aa26-27 with probability: 1.00 and cleavage probability: 0.98 resulting in a 129aa protein sequence. Signal peptide-encoding regions were removed and the resultant sequences submitted to IntFOLD-TS (Integrated Fold Recognition – Tertiary Structure) on the IntFOLD7 server (McGuffin et al., 2023, 2019) which uses an integrated trRosetta2 (Anishchenko et al., 2021) and LocalColabFold 1.0.0 (Jumper et al., 2021; Mirdita et al., 2022) to predict protein secondary and tertiary structures. Global and local model quality estimates were carried out using ModFOLD9 (McGuffin et al., 2021) and both single-chain and complex quality assessment were also conducted independently using DeepUMQA-X (Guo et al., 2022), to assess local residue accuracy (lDDT), complex interface accuracy (QS-score), and folding accuracy (TM-score).

Stereochemistry of deep-learning structural predictions was assessed using ProCheck incorporated in EMBL-EBI PDBsum (Laskowski et al., 2018). This produced a whole-model Ramachandran plot and separate Ramachandran plots by-residue for the Φ - ψ torsion angles of all residues; categorising each combination of Φ - ψ due to each residues relative position to its neighbours as either core, allowed or disallowed. Sidechain torsion angle combinations (χ_1_ - χ_2_), main-chain parameters including peptide bond planarity, Cα tetrahedral distortion, hydrogen-bond energy (ΔE_HB_) and overall G-factor were determined along with side-chain stereochemical parameters (σχ1 gauche(+), trans and gauche(-) torsion angles, pooled σχ_1_ and finally σχ_2_ trans torsion angles). To add to this analysis, main-chain bond lengths and angels, RMS distances from planarity and distorted geometry were determined.

Multimer predictions for, Mp1-Mp58, Mp1-AtVPS52, and Mp58-AtVPS52 and stoichiometric variations of combinations of these proteins were conducted using MultiFOLD (McGuffin et al., 2023) and independently with AlphaFold3 (Abramson et al., 2024) while quaternary structures were assessed using ModFOLDdock (Edmunds et al., 2023; McGuffin et al., 2023) AlphaFold3, DeepUMQA-X and ProCheck.

### M. persicae performance assays on N. benthamiana

*N. benthamiana* plants were infiltrated with pAtSUC2::Mp1 and/or pAtSUC2::Mp58 under the phloem specific AtSUC2 promoter (pHW59) (Gottwald et al., 2000). Two adult aphids were placed on the underside of leaves 1 day after agroinfiltration with clip cages. The following day adults were removed, and two 1^st^ instar nymphs were left on each infiltration site. Seven dpi, nymphs were transferred to new agroinfiltrated leaves and aphids were counted 7 days later (14 days post initial agroinfiltration).

## Acknowledgements

We thank our colleagues at the Division of Plant Sciences, University of Dundee, and in Cell and Molecular Sciences at the James Hutton Institute (JHI) for helpful discussions and advice. We would like to thank Yao Fu for help with illustrating our working model of the Mp1-Mp58-VPS52 complex. This work was supported by the European Union (ERC Consolidator grant, project number 101000997, APHIDTRAP, awarded to JIBB). The JHI Imaging Facility is funded by the Rural C Environment Science C Analytical Services Division of the Scottish Government.

## Author Contributions

JIBB conceived the study. JIBB, JRB, NG, TW, SRF designed the research and analysed the data and JRB, NG, TW, SRF performed the experiments. Specifically, JRB performed in planta Co-IPs, aphid performance assays, and confocal microscopy experiments with Mp1 and Mp58. NG performed in planta Co-IPs, as well as confocal microscopy involving Mp1, Mp58 and VPS52 with the support of MP. TW performed recombinant protein production and in vitro interaction experiments. SRF performed the computational modelling of protein (complex) structures. JRB and JIBB wrote the manuscript with input from all authors. All the authors have read and approved the paper. JRB and NG contributed equally to this work.

## Data availability statement

Data available in article and in article supplementary material.

## Supporting Information legends

**Figure S1:** co-immunoprecipitation of RFP-Mp1 and GFP-Mp58

**Figure S2:** Computational modelling of Mp1 and Mp58

**Figure S3:** Computational modelling of different Mp1-Mp58 complex conformations

**Figure S4:** AlphaFold3 and DeepUMQA-X quality analysis of the top four Mp1-Mp58 multimer predictions

**Figure S5:** PDBSum analysis of Mp1-Mp58 multimer predictions for the top four stoichiometries with the highest confidence

**Figure S6:** AlphaFold3 multimeric prediction of Mp1-AtVPS52 and Mp58-AtVPS52

**Figure S7:** PDBsum analysis of Mp1 or Mp58 and Vps52 multimer predictions

**Figure S8:** Subcellular localisation of RFP-AtVPS52, GFP-Mp1, and GFP-Mp58 expressed with GFP or RFP (controls for Fig. 3D)

**Figure S9:** Amino acid alignment of Arabidopsis VPS52 (AtVPS52) and barley VPS52 (HvVPS52)

**Figure S10:** Individual replicates of co-immunoprecipitation of RFP-VPS52 variants and GFP-Mp58

**Figure S11:** Subcellular localisation of RFP-VPS52 variants and GFP-Mp1

**Figure S12:** Subcellular localisation of RFP-AtVPS52 N-terminal deletion mutant AtVPS52^184-707^

## Supporting Information

**Figure S1:**
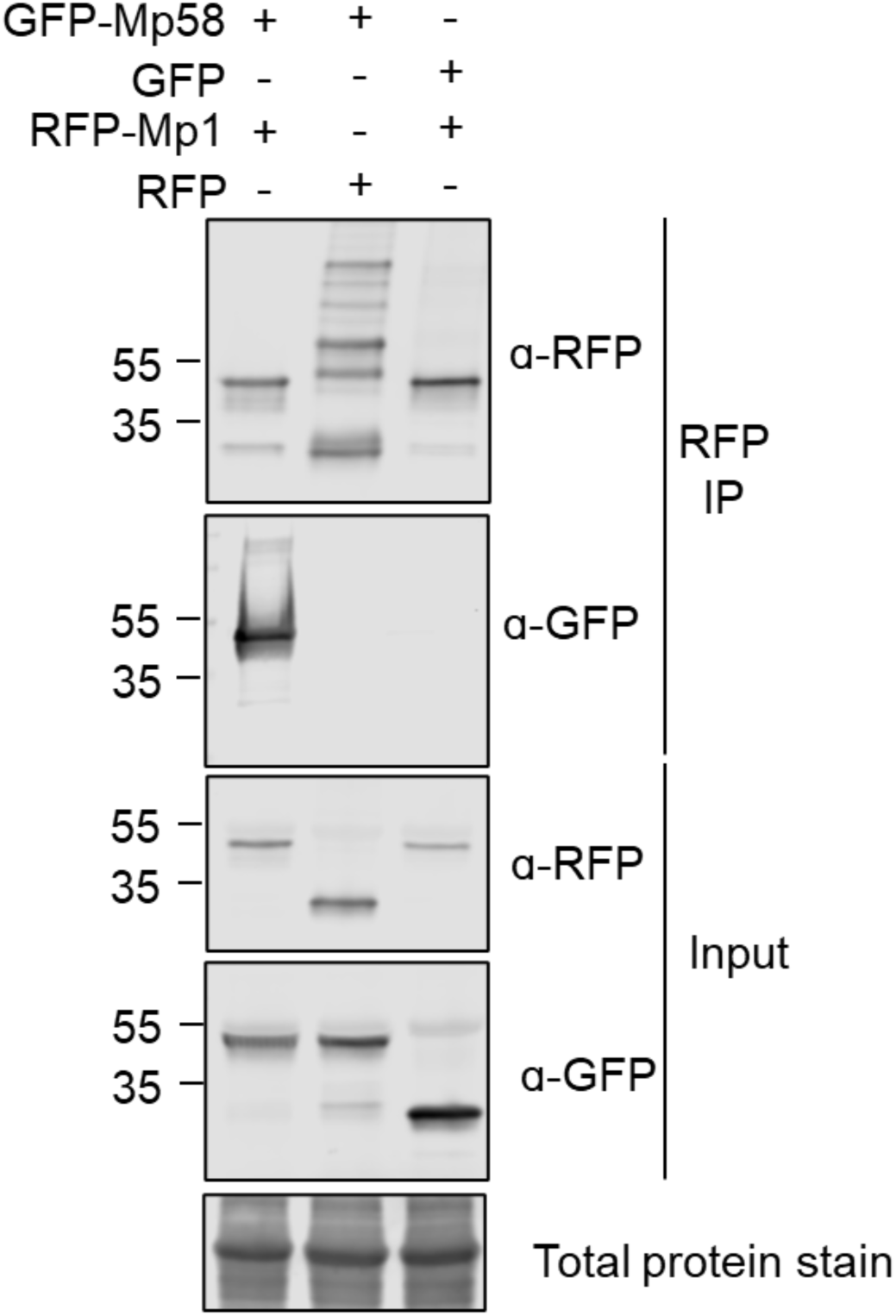
Co-immunoprecipitation of RFP-Mp1 and GFP-Mp58. RFP-Mp1 and GFP-Mp58 were transiently expressed via agroinfiltration with each other or GFP and RFP as controls. Proteins were immunoprecipitated with RFP-trap and blotted against RFP and GFP antibodies.

**Figure S2:**
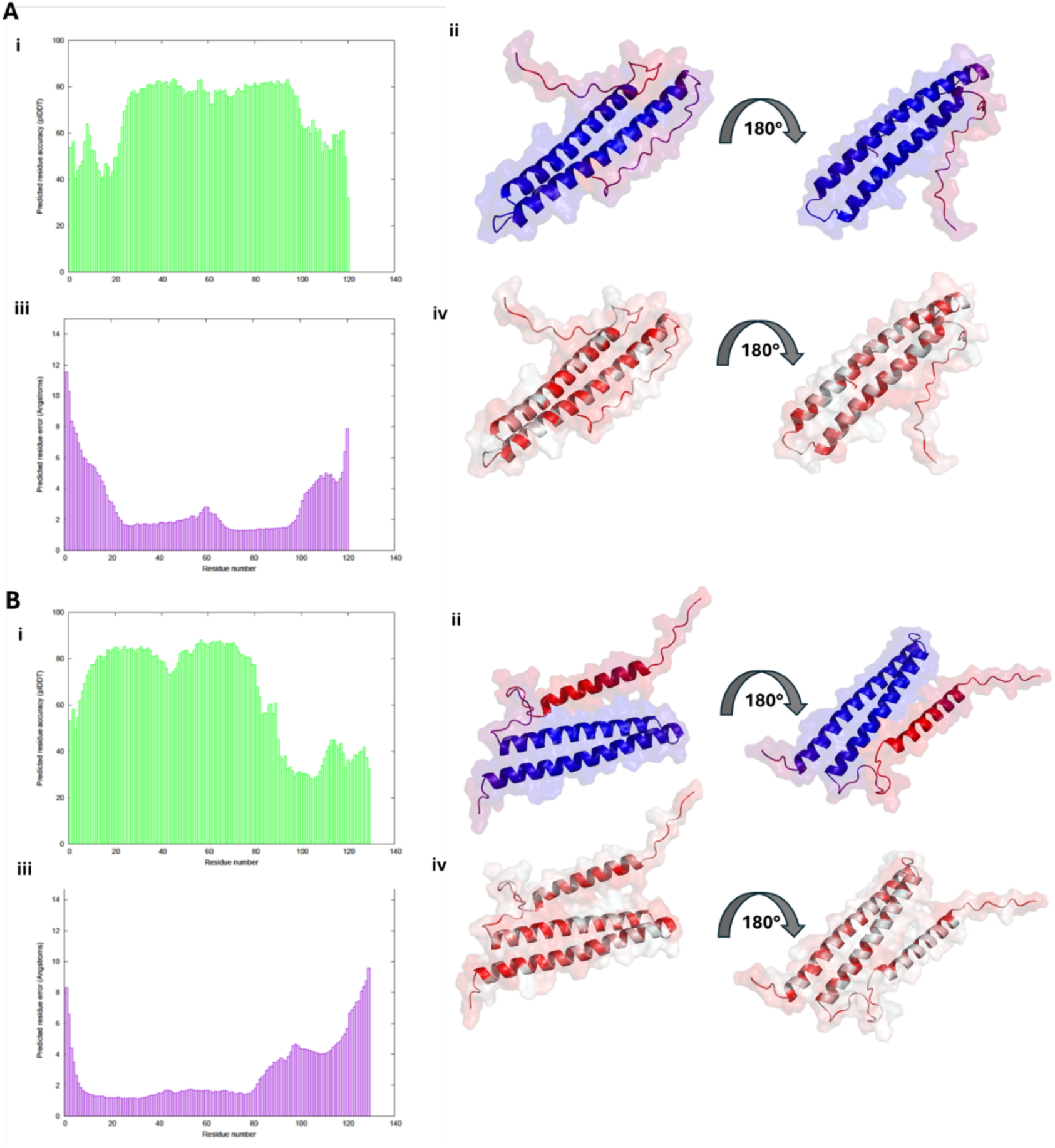
**(A)** IntFOLD7 monomeric structural prediction of Mp1 produced a model with E = 7.93e^-4^, p = 0.001, GMǪS of 0.571 and DeepUMQA-X derived global lDDT of 69.07. Figure **(i)** displays the predicted residue accuracy (plDDT) and **(ii)** shows this mapped onto Mp1 using a red-blue/low-high scale. **(iii)** displays the predicted residue error in Å. **(iv)** shows the by-residue hydrophobicity of the model, indicating intra-chain interaction and a solvent-exposed series of residues along a second side of both α-helices. **(B)** IntFOLD7 monomeric structural prediction of Mp58 produced a model with E = 1.3e^-3^, p = 0.01, GMǪS of 0.545 and DeepUMQA-X derived global lDDT of 58.86. Figure **(i)** displays the predicted residue accuracy (plDDT) and **(ii)** shows this mapped onto Mp58 using a red-blue/low-high scale. The overall plDDT is reduced due to uncertainty over the location of the Cʹ α-helix. **(iii)** displays the predicted residue error in Å. **(iv)** shows the by-residue hydrophobicity of the model, indicating intra-chain interaction between the first and second α-helices and a solvent-exposed series of residues along a second side each. The Cʹ α-helix has a single hydrophobic side.

**Figure S3:**
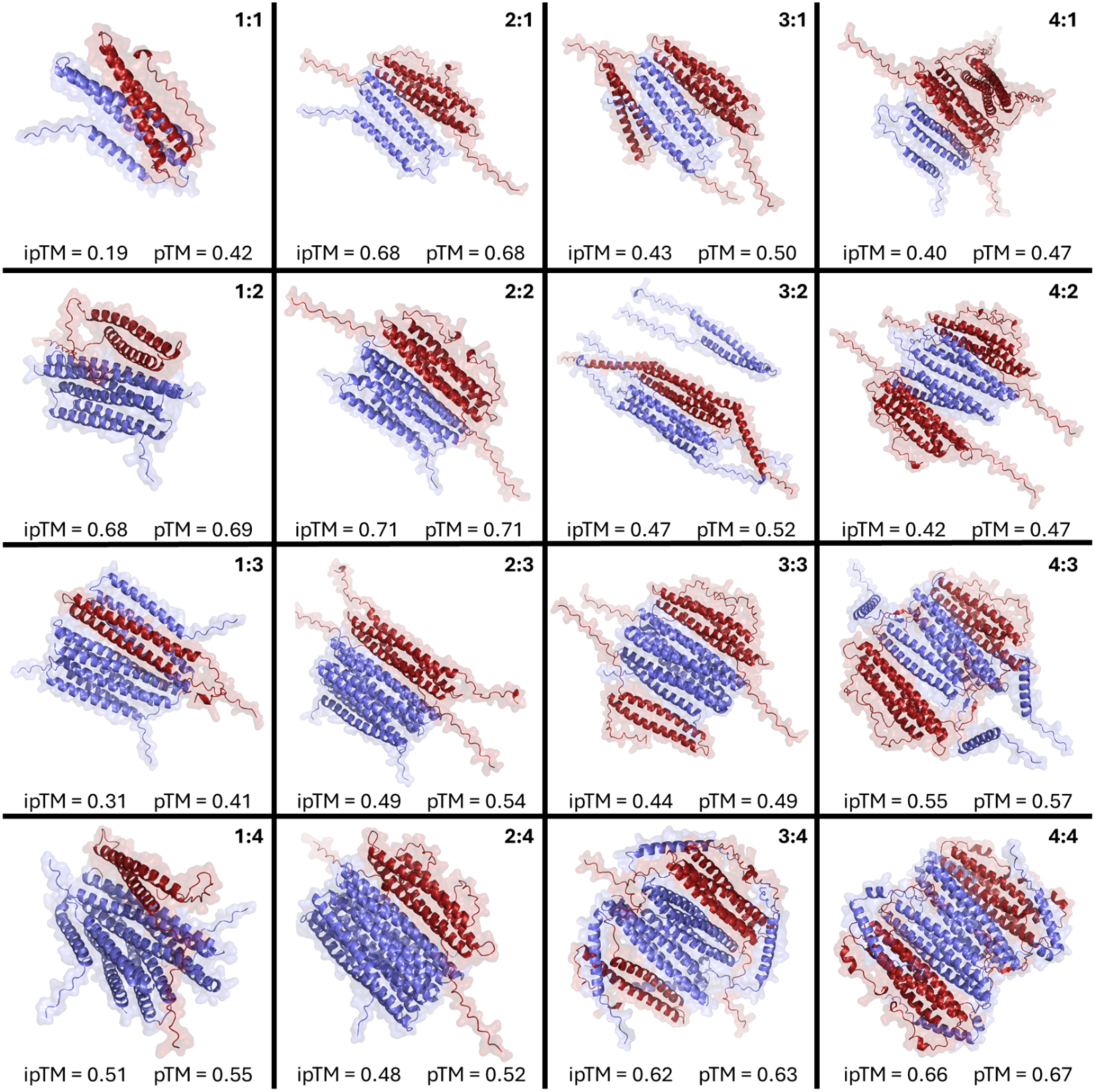
Presented are 16 possible stoichiometric permutations of the Mp1+Mp58 interaction predicted using AlphaFold3. Predicted template modelling scores (pTM) which quantify the overall accuracy of the predicted protein complex structure ranged between 0.41 and 0.71, where values > 0.5 are regarded as similar to the true structure. The interface predicted template modelling Score (ipTM) evaluates and scores the accuracy of the predicted interaction interfaces in a protein-protein complex. This ranged between 0.19 and 0.71, where > 0.8 denotes high confidence in the predicted interface, 0.6 – 0.8 moderate confidence and < 0.6 low confidence. Due to the inclusion of disordered and non-interacting domains in the input sequences used in whole-model generation, ipTM scores are consequently lower than had only the interacting domains been modelled.

**Figure S4:**
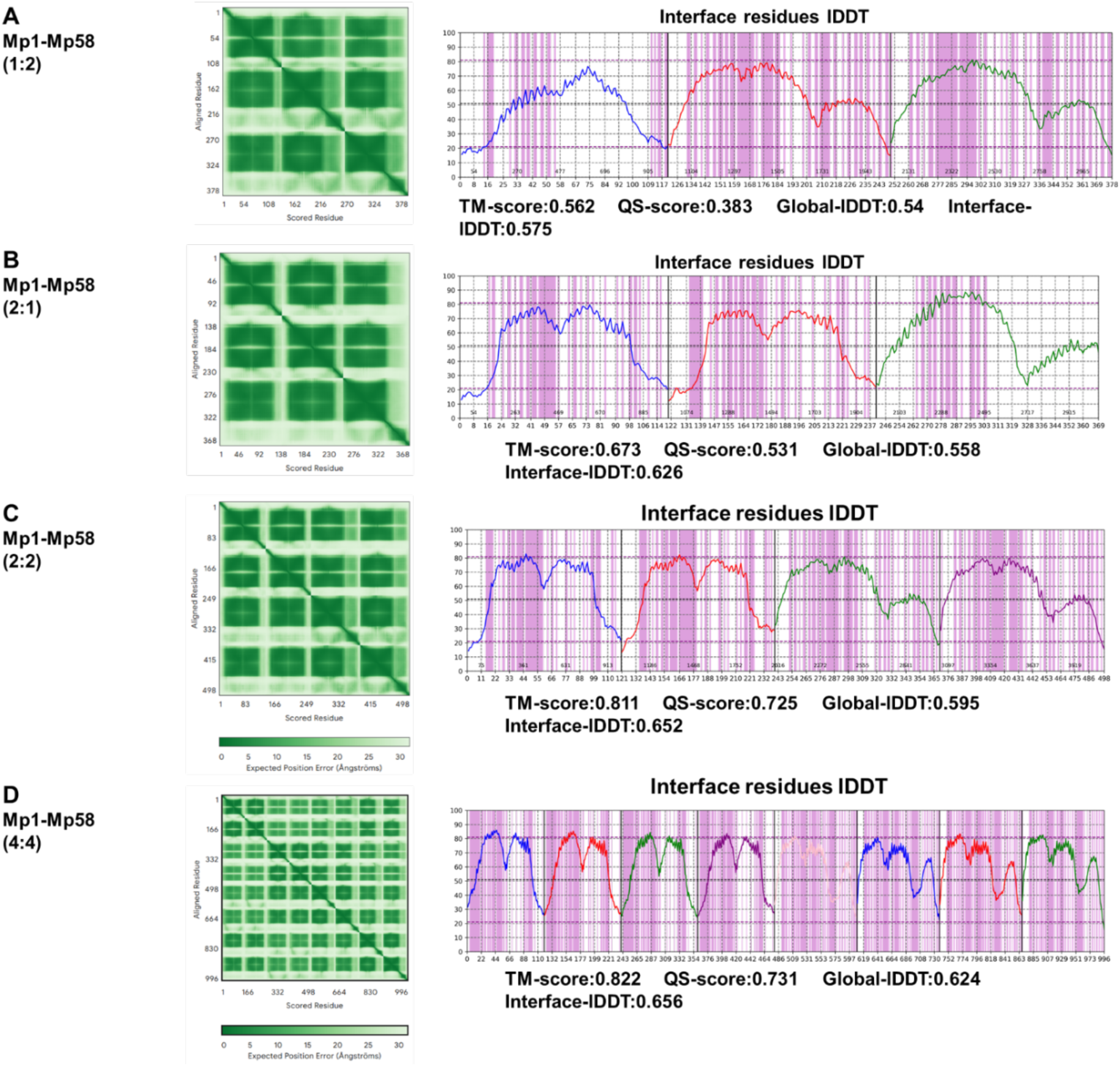
AlphaFold 3 expected position error and DeepUMQA-X model quality analysis for highest ipTM and pTM Mp1-Mp58 interaction predictions with **(A)** 1:2, **(B)** 2:1, **(C)** 2:2, and **(D)** 4:4 stoichiometries.

**Figure S5:**
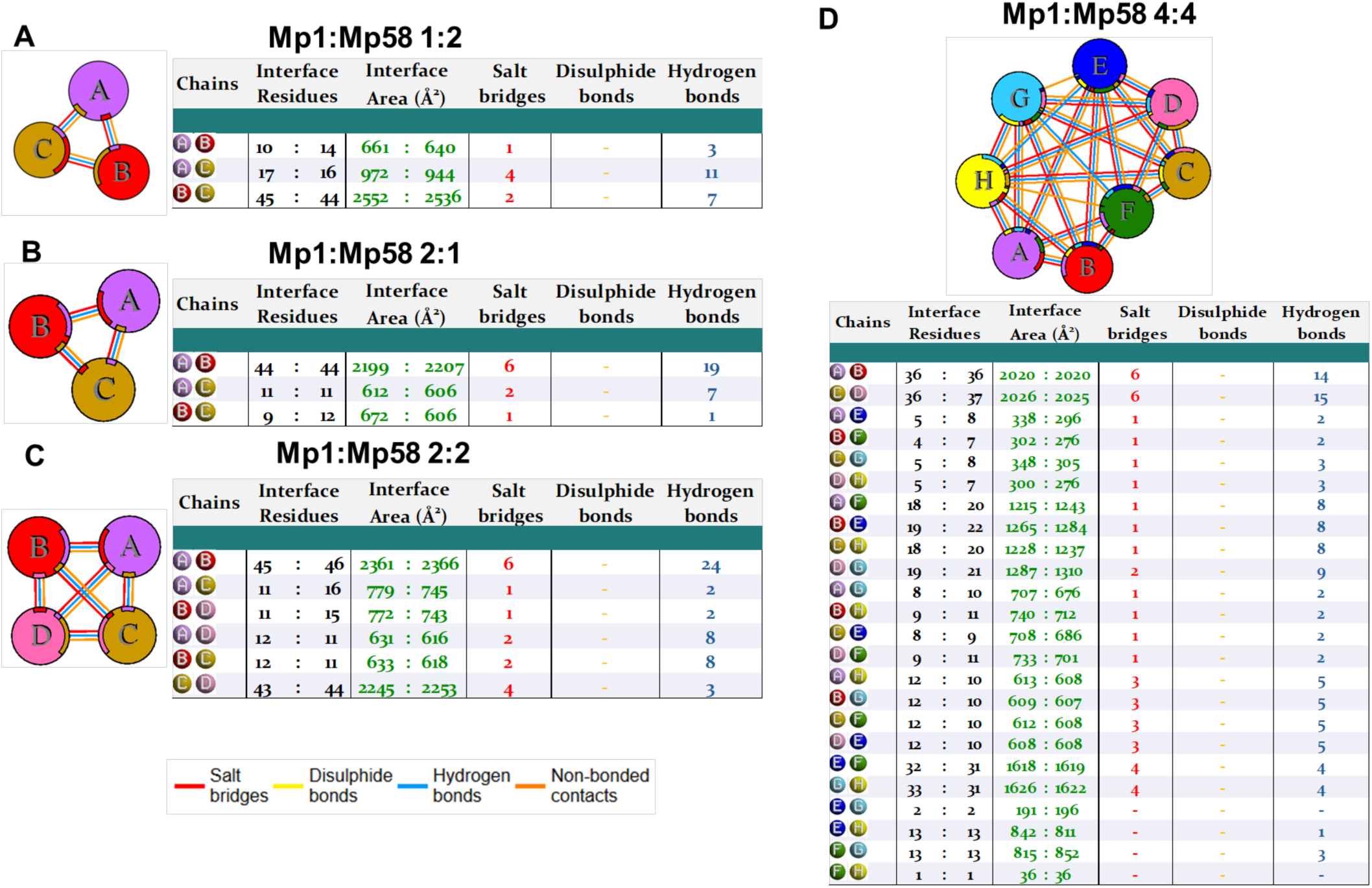
PDBsum analysis of Mp1-Mp58 multimer predictions for varying stoichiometries. Part **(A)** provides the data for 1:2 stoichiometry with chains A representing Mp1 with B and C as Mp58. **(B)** shows 2:1 with Mp1 as chains A and B while C is Mp58. **(C)** shows a 2:2 ratio with A+B as Mp1 and C+D as Mp58. Figure **(D)** shows a 4:4 ratio with A+B+C+D as Mp1 and E+F+G+H as Mp58.

**Figure S6.**
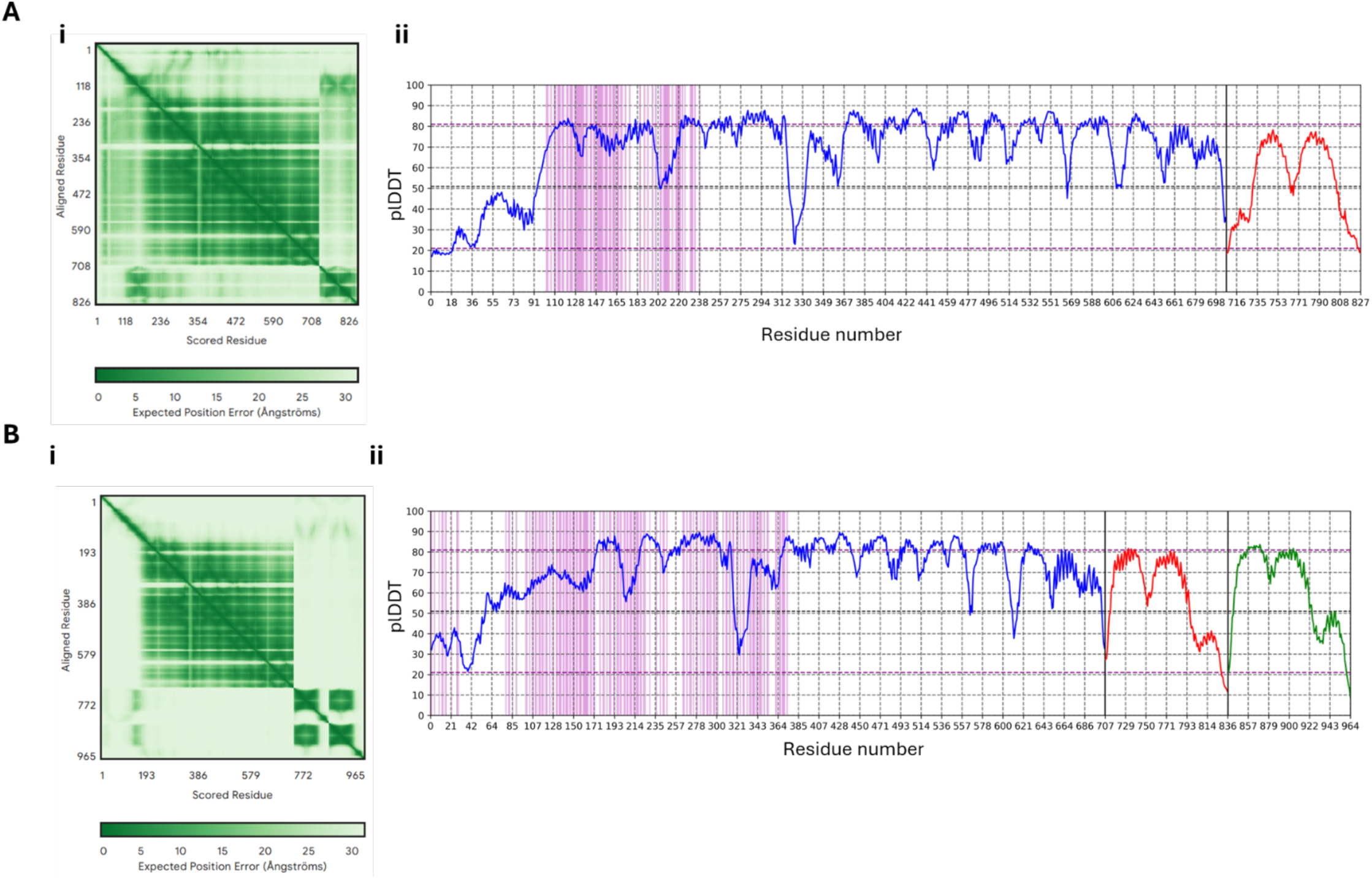
(A) AlphaFold3 multimeric prediction of Mp1-AtVPS52 with 1:1 stoichiometry produced a model where ipTM = 0.52 and pTM = 0.65. **(i)** displays the position error in Å **(ii)** DeepUMQA-X results by residue, with a whole-model TM-score: 0.816, QS-score: 0.701, Global-lDDT:0.665 Interface-lDDT: 0.632 with highlighted VPS52 contact residues in magenta **(B)** AlphaFold3 multimeric prediction of Mp58-AtVPS52 with 2:1 stoichiometry produced a model where ipTM = 0.20 and pTM = 0.52. **(i)** displays the position error in Å **(ii)** DeepUMQA-X results by residue, with a whole-model TM-score: 0.748, QS-score: 0.0.371, Global-lDDT:0.677 Interface-lDDT: 0.642 with highlighted VPS52 contact residues in magenta

**Figure S7:**
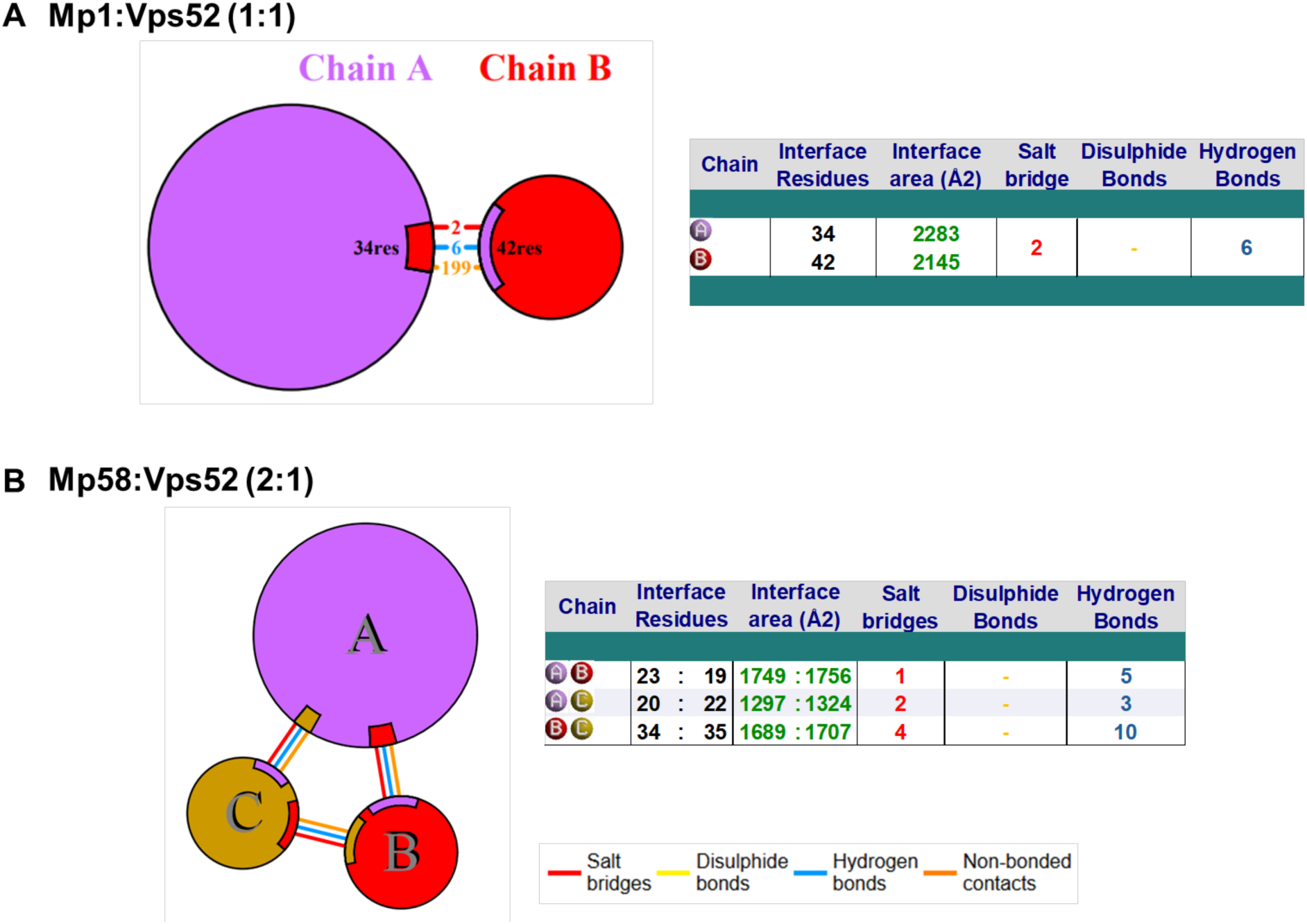
PDBsum analysis of Mp1 or Mp58 and VPS52 multimer predictions. Part **(A)** provides the data for 1:1 stoichiometry with chain A representing Vps52 and B as Mp1. **(B)** shows 2:1 with VPS52 as chain A and Mp58 as chains B and C

**Figure S8:**
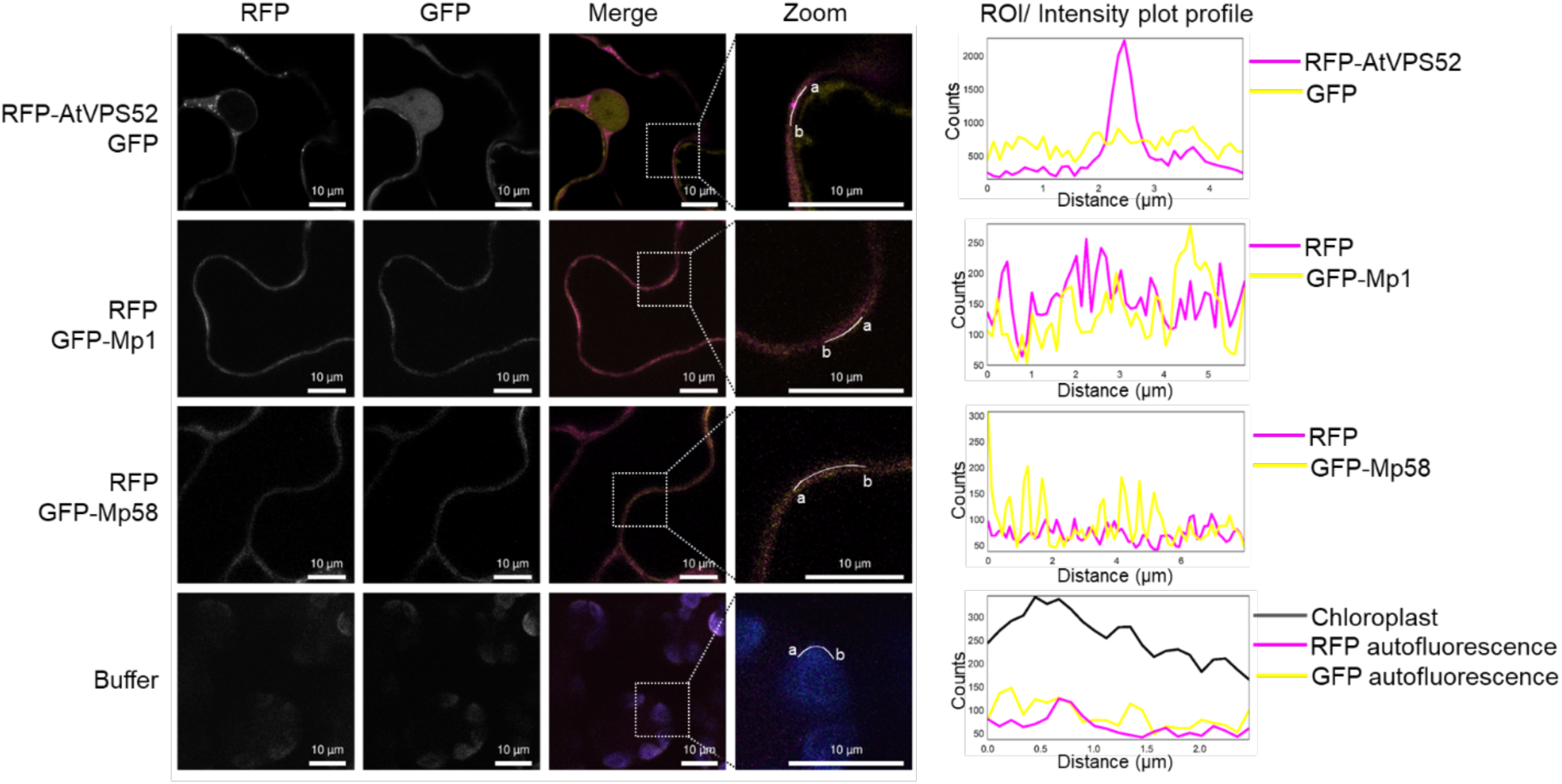
Subcellular localisation of GFP-Mp1, GFP-Mp58 (in yellow), and RFP-AtVPS52 (in magenta) (controls for Fig. 3D) Proteins were transiently expressed *N. benthamiana* via agroinfiltration and localisation was observed with confocal microscopy. Magnification = 60X (water immersion lens), scale bar = 10 µm. Presented images are single plane images. The merged panel transect correspond to line intensity plot showing fluorescence distribution across the marked locus.

**Figure S9:**
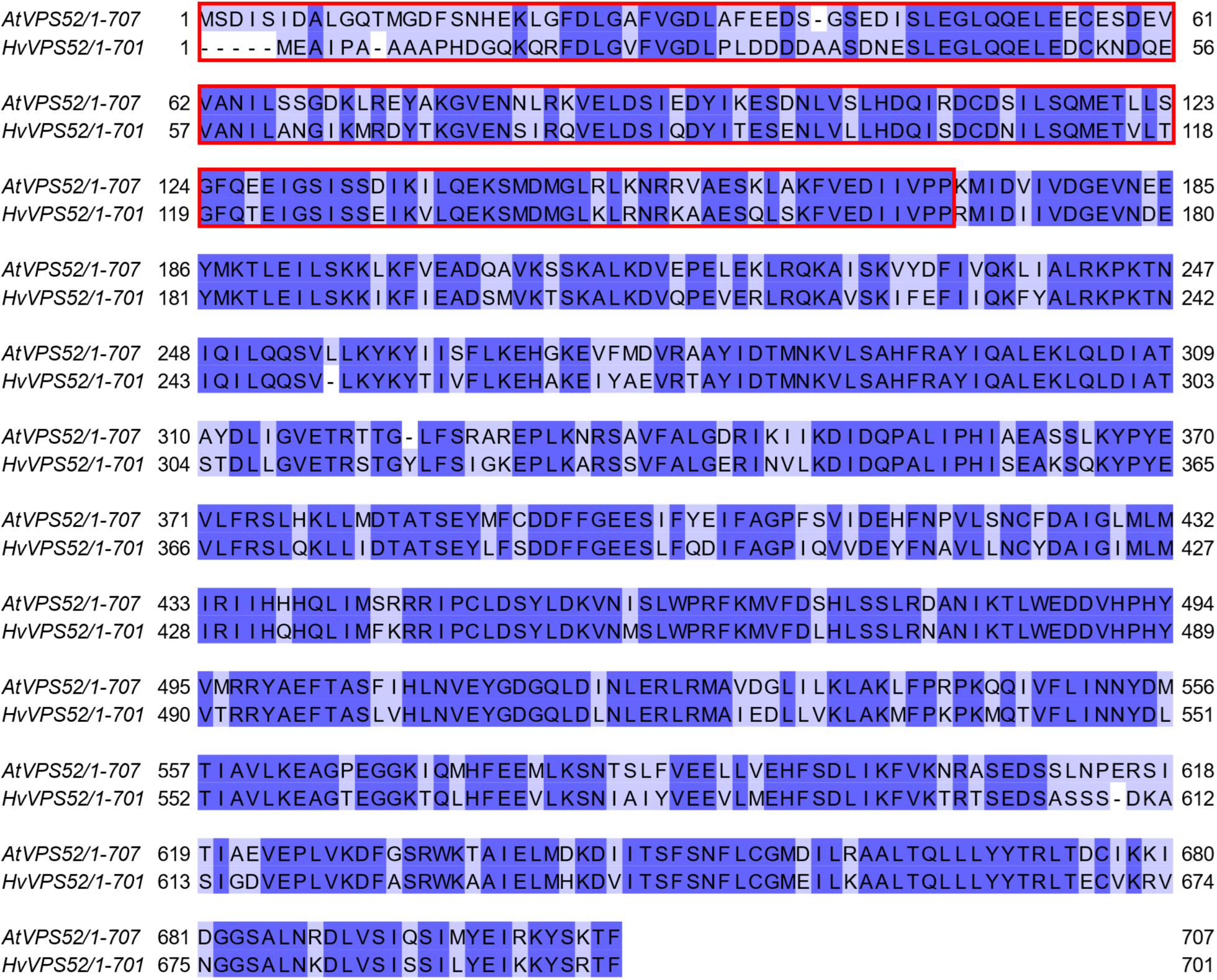
Alignment of AtVPS52 and HvVPS52 amino acid sequences. The red box indicates N-terminal region swapped in VPS52 chimeras. Alignment created in Jalview.

**Figure S10:**
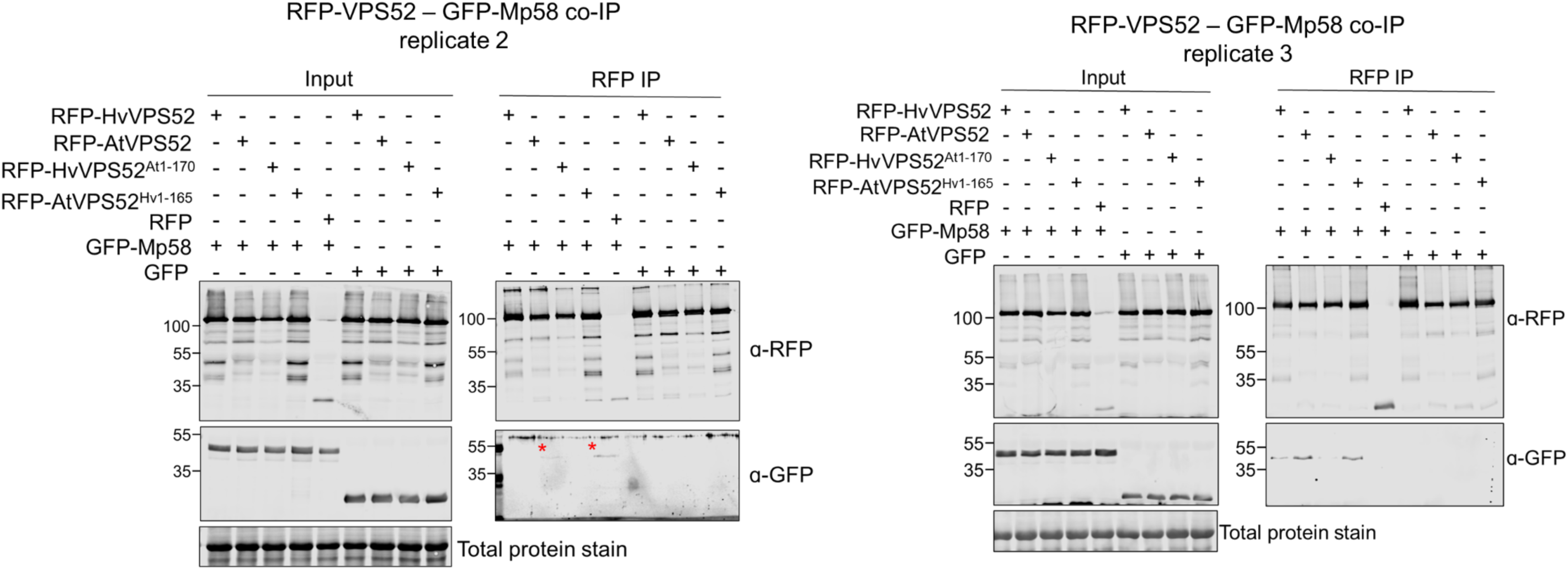
Individual replicates for co-IP of RFP-tagged VPS52 variants with GFP-Mp58 GFP-Mp58 was co-expressed with RFP-HvVPS52, RFP-AtVPS52, or N-terminal chimeras in *N. benthamiana* via agroinfiltration. RFP-tagged proteins were pulled-down with RFP-trap and blotted against GFP. RFP and GFP were used as negative controls.

**Figure S11:**
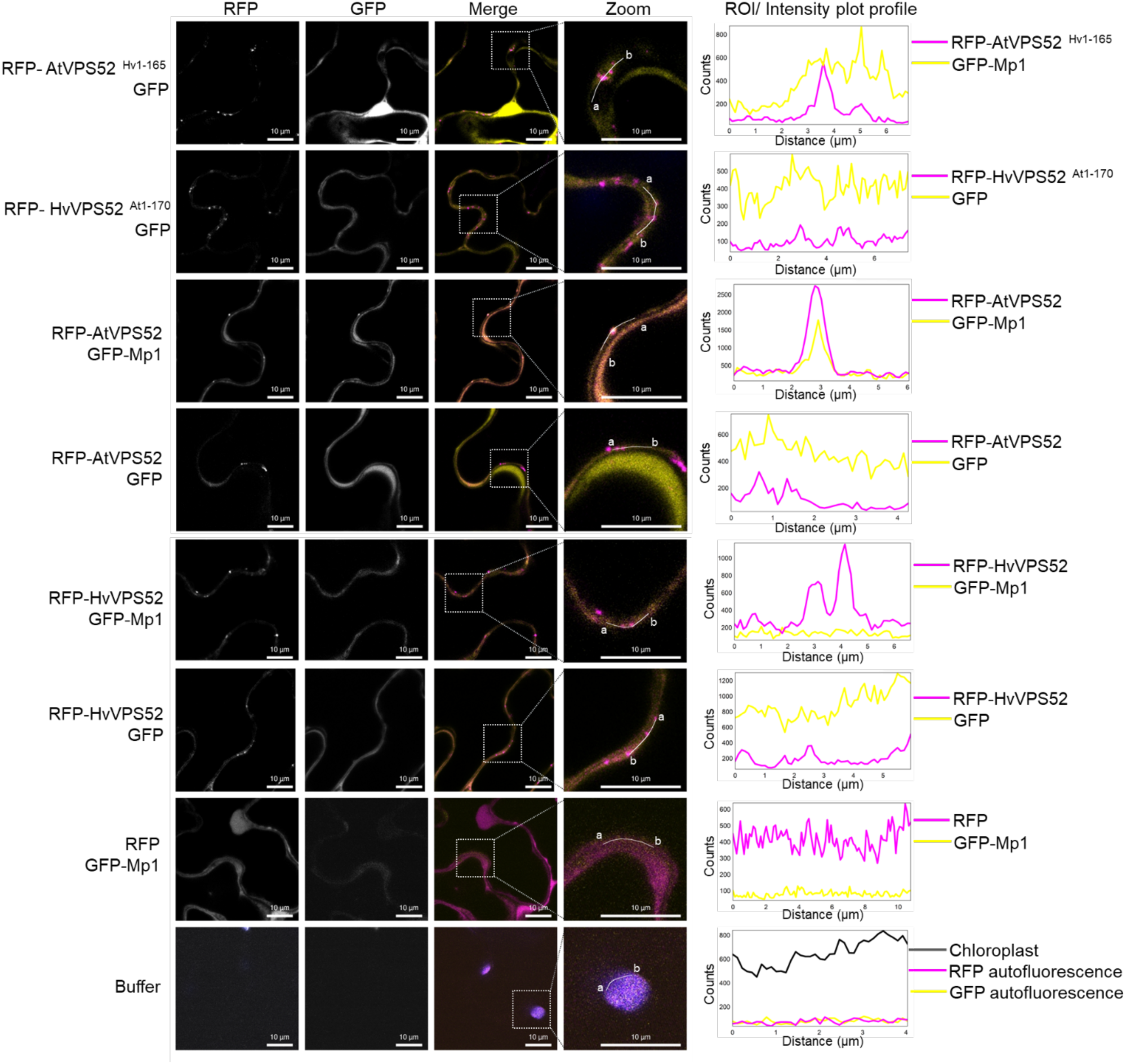
Subcellular localisation of RFP-VPS52 variants (in magenta) and GFP-Mp1 (in yellow) Proteins were transiently expressed in *N. benthamiana* via agroinfiltration, and subcellular localisation and localisation was observed with confocal microscopy. Magnification = 60X (water immersion lens), scale bar = 10 µm. Presented images are single plane images. The merged panel transect correspond to line intensity plot showing fluorescence distribution across the marked locus.

**Figure S12:**
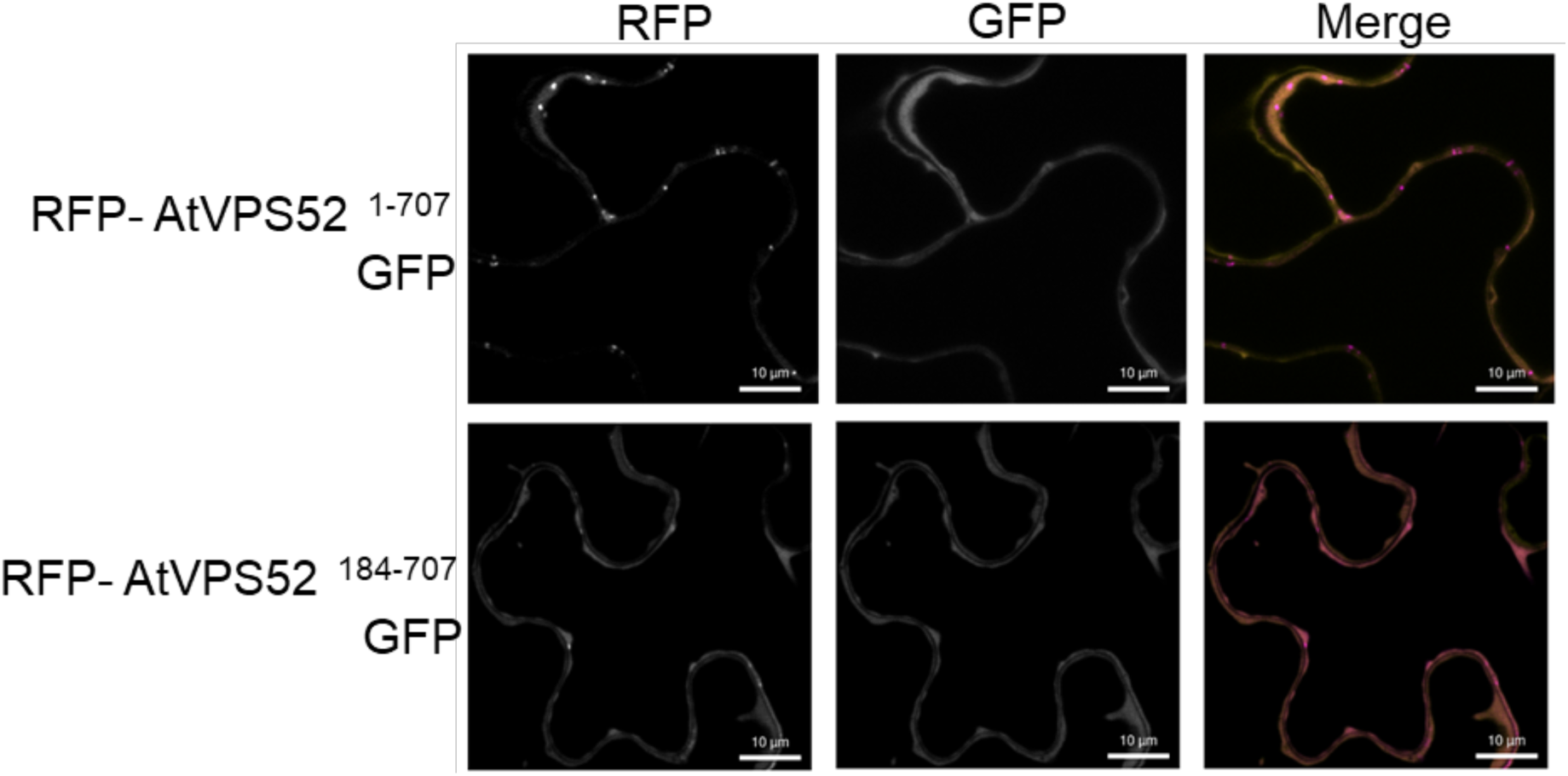
Subcellular localisation of RFP-VPS52^184-707^ (in magenta) compared to full length RFP-AtVPS52 Proteins were transiently expressed in *N. benthamiana* via agroinfiltration, and subcellular localisation and localisation was observed with confocal microscopy. Magnification = 60X (water immersion lens), scale bar = 10 µm. Presented images are single plane images. The merged panel transect correspond to line intensity plot showing fluorescence distribution across the marked locus.

**Uncropped western blots:**
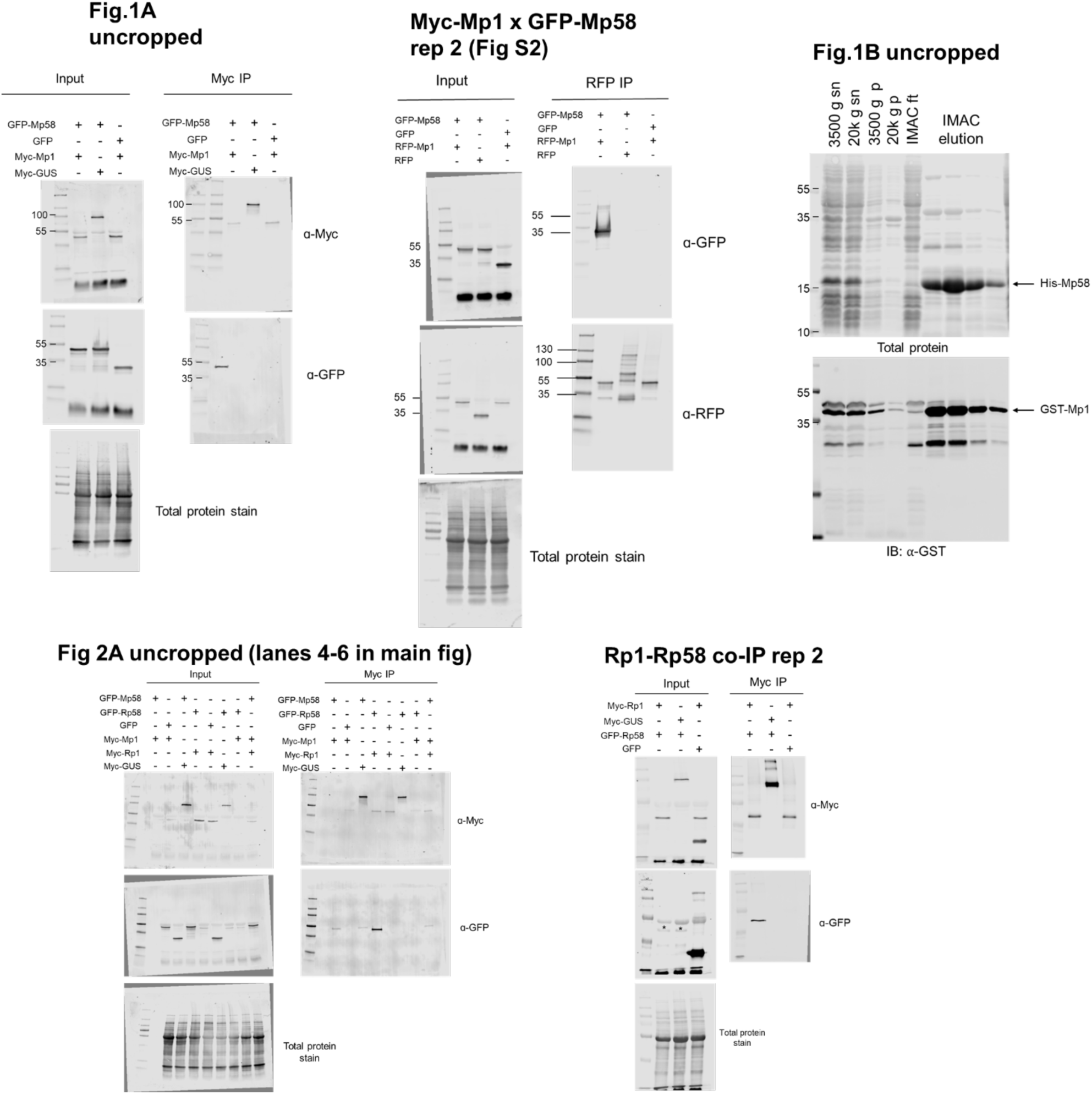
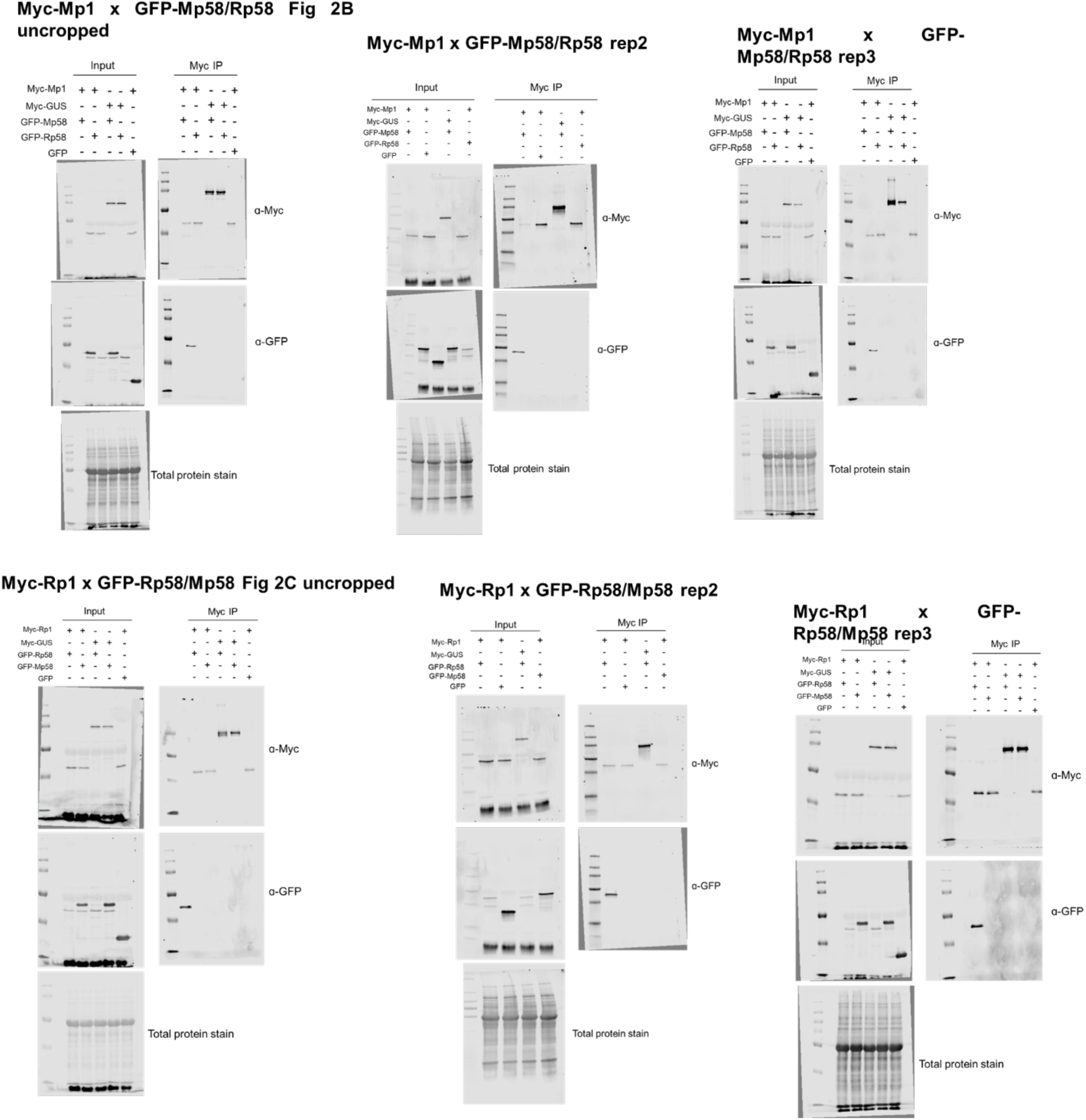
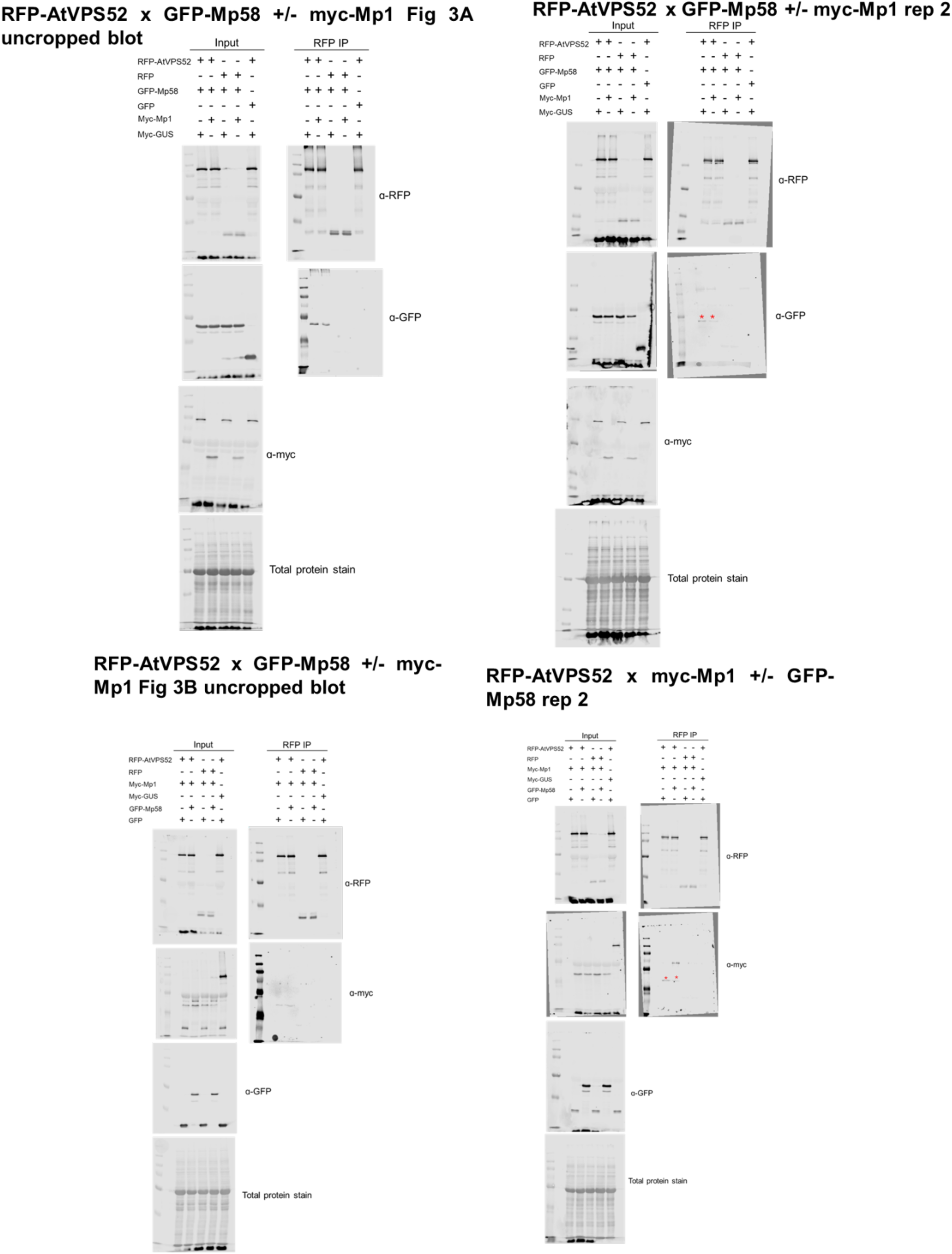
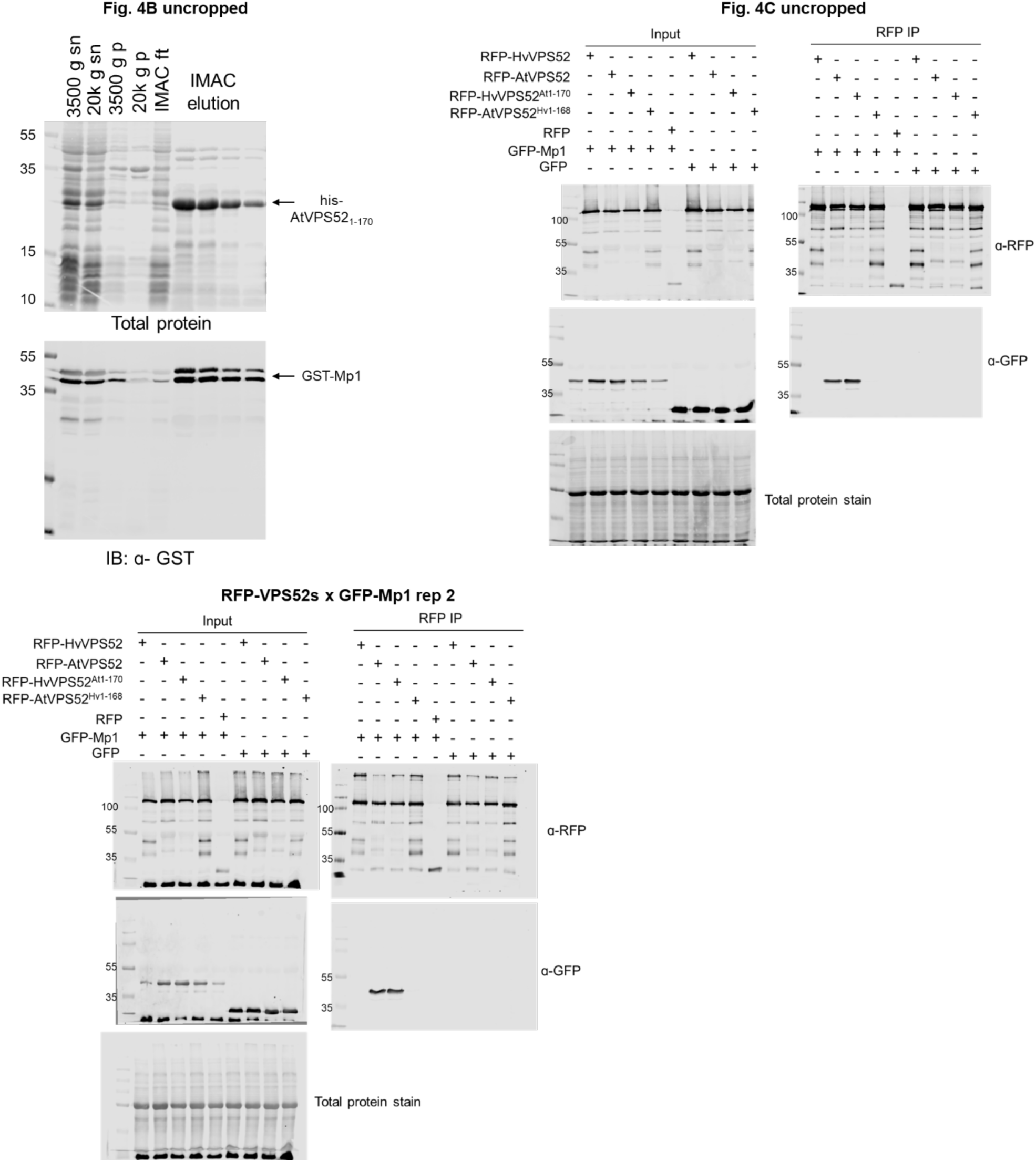
Uncropped western blots for Figs. 1, 2, 3, and 4 including additional replicates

## Notes

### Competing Interest Statement

The authors have declared no competing interest.

